# GCN5-TADA2B in the SAGA complex provide constitutive fine-tuning control of XRCC1 recruitment and focal retention at DNA damage sites

**DOI:** 10.1101/2025.08.24.672030

**Authors:** Keeeun Kim, Junyoung Kim, Darom Lee, Jae-Hoon Ji, Myung-Hee Kwon, Byong Chul Yoo, Youngsoo Lee

## Abstract

The scaffolding protein XRCC1 orchestrates base excision repair (BER) and DNA single strand break repair (SSBR) through extensive protein-protein interactions, and its disruption contributes to human neurological diseases such as SCAR26. Recruitment to DNA damage sites has been traditionally understood to depend exclusively on interaction with PARP1 and PARP1 mediated PARylation. Here, we report for the first time the discovery of GCN5 and TADA2B, components of histone acetyltransferase module of SAGA complex, as novel XRCC1 binding partners that provide critical fine-tuning control over repair site localization. We demonstrate that these SAGA complex components bind XRCC1 constitutively via distinct BRCT domain interactions — GCN5 mainly to BRCT I and TADA2B mainly to BRCT II — independent of DNA damage status or GCN5’s acetyltransferase activity. Depletion of either protein significantly impairs XRCC1 recruitment efficiency, while paradoxically rescuing focal retention defects in BRCT II deficient XRCC1 mutants. This dual regulatory behavior reveals a regulatory mechanism where constitutive protein interactions optimize normal XRCC1 function but become counterproductive when XRCC1 is structurally compromised, particularly in the BRCT II domain. Analysis of the SCAR26 associated variant, which disrupts TADA2B binding while maintaining LIG3 interaction, confirms this dual regulatory model and provides molecular insights into potential disease pathophysiology. Our findings establish a ‘ready-for-action’ model of DNA repair coordination where protein complexes pre-organize repair machinery for immediate deployment upon damage detection, expanding our understanding of BER regulation.

## INTRODUCTION

The nervous system is the most complex organ in the body controlled by various regulatory mechanisms, including DNA damage response (DDR) and repair, which play crucial roles in proper development and function of the nervous system ^1,2^. Multiple animal models with defects in DNA repair mechanisms have indicated that endogenous DNA damage or strand breaks naturally occur during brain development, although the precise nature of this spontaneous DNA damage remains incompletely understood ^3–11^. Consequently, impaired responses to endogenous DNA damage in the nervous system can lead to abnormal neurodevelopment and neurological disorders ^2,12^.

Ataxia telangiectasia (A-T) represents one of the most studied rare genetic diseases in this physiological context. Patients with A-T exhibit progressive cerebellar ataxia accompanied by several non-neural symptoms, including immune deficiency, radiosensitivity, and tumor susceptibility ^13^. This disease is caused by mutations in the Ataxia Telangiectasia Mutated (*ATM*) gene, which encodes a protein kinase that phosphorylates numerous substrates, including ATM itself, H2AX, KAP1, CHK2, p53 to regulate damage repair, DDR, cell cycle progression, and programmed cell death in response to DNA damage, particularly DNA strand breaks ^14–16^. The instant recruitment of ATM to DNA damage sites depends on the MRE11-RAD50-NBS1 (MRN) complex, and mutations of MRE11 result in A-T like disorder (ATLD), which presents with cerebellar ataxia in a slower progression than classical A-T ^12,14^. While the direct connection between neurological phenotypes, particularly cerebellar dysfunction, and DNA repair defects is evident, the underlying molecular mechanisms are not fully elucidated.

Furthermore, the list of human genetic diseases with neurological phenotypes linked to DNA repair defects continues to expand. This includes spinocerebellar ataxia with axonal neuropathy1 (SCAN1, caused by mutations in TDP1), ataxia oculomotor apraxia 1/4 (AOA1/AOA4, caused by mutations in APTX and PNKP respectively), and microcephaly with seizure (MCSZ, caused by mutations in PNKP). These disorders result from defective DNA base excision repair (BER) or single strand break repair (SSBR) mechanisms ^12,17^. Notably, all these proteins, despite their diverse roles in BER or SSBR, are recruited to DNA damage sites in an XRCC1 dependent manner through specific protein-protein interactions ^18,19^.

To investigate the connection between BER/SSBR mechanisms and neurodevelopment/neural function, the *Xrcc1* conditional knockout animal model has been generated and analyzed. This animal model, in which the *Xrcc1* gene is deficient in the nervous system, exhibited spontaneous seizure, mild ataxia, and notably, interneuron loss in the cerebellum due to cell cycle arrest ^20^. Subsequently, a human case with compound heterozygous mutations in the *XRCC1* gene was reported, presenting with adult-onset spinocerebellar ataxia, progressive cerebellar atrophy, and axonal neuropathy, which is now named SCAR26 (Spinocerebellar ataxia, autosomal recessive 26) ^21^. Considering XRCC1 functions as a scaffolding protein that binds multiple DNA repair proteins and facilitates their recruitment to DNA damage sites ^18,19^, the neurological phenotypes observed in the *Xrcc1* conditional knockout model likely result from the absence of one or more XRCC1 binding partners at sites of DNA damage. Intriguingly, none of the animal models with deficiencies in known *Xrcc1* binding partners including DNA polymerase β (*Polb*), DNA ligase III (*Lig3*), *Tdp1*, *Pnkp*, and *Aptx* reproduced the cerebellar phenotypes observed in the *Xrcc1* deficient animals. Each animal model of XRCC1 binding partners showed distinct neurological phenotypes ^6,8–11,22^. This suggests that other XRCC1 interacting proteins, possibly yet undiscovered, might be responsible for these specific cerebellar manifestations observed in the *Xrcc1* animals.

As it relates to that, we identified TADA2B (Transcriptional adaptor 2B) and GCN5 (General Control of Non-depressible 5, KAT2A - lysine (K) acetyltransferase 2A) as novel XRCC1 binding proteins. TADA2B is a component of the SAGA (Spt-Ada-Gcn5 acetyltransferase) histone acetyltransferase (HAT) module ^23–25^. The interaction between TADA2B and GCN5 is required for the full HAT enzyme activity of GCN5 ^24,26^. Furthermore, the SAGA complex, particularly TADA2B, plays an important role in pluripotency, survival, growth, and lineage specification in embryonic stem cells. TADA2B mutants confer a selective advantage in human embryonic stem cells, but not in somatic cells ^27^. Although ADA2b in *Arabidopsis* (homologous to human TADA2B) is localized to DNA double strand break (DSB) sites and influences DNA repair ^28^, the specific role of TADA2B in DNA damage repair remains unexplored in higher organisms, including humans. On the other hand, the primary function of GCN5 is to regulate gene activation through histone acetylation, particularly histone H3K9 and H3K14, as well as acetylation of numerous non-histone targets including DNA repair proteins such as K830 of BRCA1, K36/37 of PARP2, and K177 of WEE1 to enhance their functions in DNA repair and responses ^29–31^. Upon UV induced DNA damage, GCN5 mediated acetylation of RPA1 at K163 augments DNA damage repair via the protein interaction with XPA, one of key factors in nucleotide excision repair (NER), while the RPA complex protects the complementary DNA strand opposite to the damaged strand ^29,32,33^. Some of these functions and target protein modifications are shared with PCAF (p300/CBP associated factor, also known as KAT2B) which, unlike GCN5, does not bind to TADA2B ^29,30^. Furthermore, GCN5 exhibits additional acyltransferase activities, including propionylation and butyrylation of lysine residues on histones, which have been observed to regulate transcription ^34,35^. Overexpression of GCN5, found in acute myeloid leukemia (AML) patients with poor survival, enhances ATM activation in response to DNA damage in early drug resistant leukemia cells r^36^. Loss of *Gcn5* in the animal model causes early embryonic lethality, while the murine model with point mutations in the acetyltransferase domain of GCN5 showed developmental defects in the nervous system, including cranial neural tube closure defects and exencephaly, as well as abnormalities in neural crest cell derived non-neuronal organs ^30^. A *Tada2b* murine model has not yet been generated.

In the current study, we identify GCN5 and TADA2B as new binding partners of XRCC1 and demonstrate that GCN5 and TADA2B are necessary for the precise orchestration of XRCC1 recruitment to DNA damage sites. While XRCC1 recruitment fundamentally depends on PARP1 activation and target protein PARylation (Poly ADP-ribosylation) at damage sites ^18,37^, we discovered that GCN5 and TADA2B serve as previously unidentified facilitators in this recruitment and focal retention of XRCC1. Although we do not have an answer to whether GCN5 and/or TADA2B are responsible for neuronal phenotypes observed in *Xrcc1* deficient cerebellum, our findings reveal that XRCC1 interaction with GCN5/TADA2B under normal cellular conditions likely pre-positions XRCC1 in proximity to DNA, thereby enabling its rapid mobilization to and maintenance of focal accumulation at DNA damage sites for efficient repair. This newly recognized mechanism represents another critical preparatory regulatory mechanism for proper and precise DNA repair mechanisms in BER and SSBR.

## MATERIALS AND METHODS

### Animal

The animal model in which the *Xrcc1* gene was selectively deleted in the nervous system by breeding with the *Nestin-Cre* animal model (*Xrcc1^Nes-Cre^*) has been described before ^11,20^. Embryonic and mature brains were removed at the indicated time points and prepared for GST (Glutathione S-Transferase) pulldown and mass spectrometry analysis. All animals were housed in the Laboratory Animal Research Center of Ajou University Medical Center. Animal usage for this study was in accordance with the guidelines from the Institutional Animal Care and Use Committee.

### Protein Mass spectrometry

To identify XRCC1 binding proteins, the murine *Xrcc1* gene was cloned into a GST-tagged vector using the Gateway cloning system (Invitrogen) (detailed information in the gene cloning section below). Glutathione Sepharose 4B (GST-beads, Cytiva) was packed into a multipurpose mini spin column (BioVision) and incubated with GST-mouse XRCC1, produced in Sf9 cell line, of which lysate was prepared in 50 mM Tris pH 8.0, 0.15 M NaCl, 1% NP-40, 1 mM phenylmethylsulfonyl fluoride (PMSF), 1 μg/ml aprotinin, and 1 μg/ml leupeptin. After multiple washing steps, wildtype (WT) and *Xrcc1* null (*Xrcc1^Nes-Cre^*) mouse brain tissue lysates, precleared with GST-beads, were applied to the GST-mouse XRCC1 column. The final elutes from the column were subjected to protein mass spectrometry analysis.

Protein mass spectrometry was performed as previously described ^38^. Briefly, Proteins in the final eluates were separated by SDS-PAGE, then followed by Colloidal blue staining (Suppl. Fig. 1A). The protein bands were also checked by protein silver staining (Dodeca Silver stain kit, Bio-Rad Laboratories). The PAGE gels were sliced into several pieces guided by blue staining and subjected to tryptic digested using an in-gel tryptic digestion kit (Thermo Fisher Scientific) according to the manufacturer’s protocol. The alkylated final gel pieces were dehydrated in acetonitrile and digested with high-grade trypsin for 12 hours at 30 °C. The tryptic digested peptides were analyzed using the Q Exactive hybrid quadrupole-orbitrap mass spectrometer with an Ultimate 3000 RSLCnano system (Thermo Fisher Scientific). All mass spectrometry samples were analyzed using Sequest (v.27, Thermo Fisher Scientific). This analysis program matched the outcome from the spectrometry with the Uniprot_sprot database using a parameter of a fragment ion mass tolerance of 1 Da and a parent ion tolerance of 1.2 Da.

### Gene cloning

cDNA was synthesized using RNA templates extracted from mouse postnatal day 5 cerebellum or commercially available human brain total RNA (Ambion) according to the manufacturer’s instructions (Invitrogen). The mouse *Xrcc1*, and human *GCN5* and *TADA2B* genes were amplified from the cDNA via PCR using primer pairs shown in supplementary (Suppl.) table 1. The human *XRCC1* gene was amplified from pGEX-h*XRCC1* (Addgene) using the PCR primer pair (Suppl. Table 1). The PCR product of each gene was cloned into an Entry vector (pCR^TM^8/GW/TOPO), and subsequently sub-cloned into GST-tagged (inducible pDEST^TM^15, constitutive pDEST^TM^27 or baculovirus expression pDEST^TM^20 system), GFP-tagged (pcDNA^TM^-DEST53) or mCherry-tagged (362 pCS Cherry DEST) vectors using the Gateway cloning system (Invitrogen). Using the cloned Entry vectors as templates, truncated mutants of each gene were amplified with the designated PCR primer pairs (Suppl. Table 1) and cloned into a GST-tagged Gateway vector. Additionally, an engineered SFB (S protein tag, Flag tag / Streptavidin binding peptide) vector with pIRES2-EGFP backbone (Takara Bio) was used for further cloning of each gene.

In addition, three single nucleotide missense mutations of the *XRCC1* gene related to SCAR26 have been selected (https://www.ncbi.nlm.nih.gov/clinvar/) ^21^. There mutations are c.1196A>G (p.Gln399Arg), c.1293G>C (p.Lys431Asn), and c.1738C>T (p.Arg580Trp), which are located in the BRCT I, the CTL, and the BRCT II domains respectively. Using the GFP-XRCC1 construct as a template, each point-mutated construct was amplified by PCR using the primer pairs indicated in Suppl. Table 1 with Pfu Turbo (Agilent). The PCR products were treated with DpnI restriction enzyme (New England Biolabs) to remove any methylated DNA strands, then cloned into GFP, SFB or GST tagged vectors. The intended point mutations were confirmed by whole gene sequencing of the *XRCC1* constructs (Suppl. Fig. 10B).

### Cell culture, gene overexpression, gene knockdown

For *in vitro* analyses, 293T/293TN (immortalized human embryonic kidney cell), U2OS (human bone osteosarcoma epithelial cell), and HeLa (human cervical cancer cell) cell lines were maintained in DMEM (high glucose with pyruvate, Gibco) supplemented with 10% fetal bovine serum (FBS, Gibco) and antibiotic-antimycotic solution (AA, Gibco) at 37 °C with 5% CO_2_. Sf9 (insect ovarian epithelial cell) cell line was maintained in Sf-900 II SFM (serum-free medium, Gibco) supplemented with 10% FBS and AA in a 28 °C incubator (Coretech).

Mouse XRCC1 protein was produced as GST-tagged protein using the baculovirus expression vector system in Sf9 cell line. The gene construct was mixed with Cellfectin II reagent (according to the user guide from Gibco) to transfect the insect cell line, followed by viral transduction of the same cell line to boost the protein production. In addition, GST-tagged human proteins were generated in an Isopropyl-β-D-thiogalactoside (IPTG, 400 μM, Sigma-Aldrich) inducible Gateway system. To overexpress the gene of interest, target cells were transfected with the cloned gene using polyethyleneimine (PEI, 1 mg/ml, Polysciences) and Opti-MEM (Gibco) for more than 12 hours. To knock down the expression of the target gene, siRNA (small interfering RNA) sequences were designed using the public websites as indicated in Suppl. table 2-1. Cells were incubated in a mixture of siRNA primers (scrambled primers for the control groups), Lipofectamine RNAiMAX (according to the transfection protocol from Invitrogen) and Opti-MEM for more than 16 hours. Transfected cells were analyzed or treated with DNA damage inducing drugs 48 to 72 hours after transfection. When needed, MG132 (10 μM, Selleckchem) was added to the culture media for 12 hours. DNA damage to cell lines was induced chemically by treatments with methyl methanesulfonate (MMS, Sigma-Aldrich), H_2_O_2_ (Sigma-Aldrich), Phleomycin (Phleo, Sigma-Aldrich) or Camptothecin (CPT, Thermo Fisher Scientific). The doses and time points for DNA damage induction are indicated in the figures. Additionally, Butyrolactone 3 (MB-3, GCN5 inhibitor, 100 μM working concentration, Cayman Chemical) was applied to the cells 1 hour prior to DNA damage induction.

### Generation of *XRCC1* knockout cell line

To generate a *XRCC1* knockout cell line, the CRISPR (Clustered Regularly Interspaced Short Palindromic Repeats)–Cas (CRISPR-associated) 9 methodology was applied to the U2OS cell line ^39^. The oligomer to target the XRCC1 gene was designed using the publicly available CRISPR-Cas9 design websites (Suppl. Table 2-2). The guide RNA for the *XRCC1* gene was cloned into the pXPR_001 vector (LentiCRISPR, GeCKO). For the packaging process of the targeted lentivirus, pMD2.G (Addgene) and psPAX2 (Addgene) were co-transfected with the cloned LentiCRISPR into 293TN cells. Next, U2OS cells were infected with media containing the targeted lentivirus from 293TN culture in combination with polybrene (Millipore, 5 μg/ml) to increase infection efficiency. *XRCC1* lentivirus-infected U2OS cells were distributed in 96-well plates, and single cell colonies were selected and examined for XRCC1 deficiency by Western blot analysis.

### Immunoprecipitation and Western blot

The antibodies used for Western blot analysis were listed in Suppl. Table 3. Mouse brain tissues were collected at the indicated time points, and processed as reported previously ^11,20^. Briefly, brain tissues were prepared in a lysis buffer containing 50 mM Tris buffer pH 7.5, 150 mM NaCl, 50 mM NaF, 0.2% NP-40, 1% Tween-20, 1 mM 4-(2-Aminoethyl)benzenesulfonyl fluoride hydrochloride (AEBSF, Sigma-Aldrich), 1 mM dithiothreitol (DTT), protease inhibitor (Sigma-Aldrich), and phosphatase inhibitor (Sigma-Aldrich). Cultured cells on plates were processed directly in a lysis buffer containing 60 mM Tris buffer pH 6.8, 2.4% sodium dodecyl sulfate (SDS), 6% β-mercaptoethanol, 0.12% bromophenol blue, and 12% glycerol for routine Western blot analysis.

IPTG-induced bacterial cells were lysed in CelLytic B Cell Lysis reagent (Sigma-Aldrich) supplemented with 50 μg/ml lysozyme, 10 μg/ml DNase, 1 mM PMSF, 1 μg/ml aprotinin, 1 μg/ml leupeptin, and 0.2% Triton X-100 in PBS, followed by incubation with GST-beads for pulldown experiments. GST-beads were washed with buffer containing 500 mM NaCl, 1 mM EDTA, 10 % glycerol, 0.1% Triton X-100, 0.07% βmercaptoethanol, and 1 mM PMSF in a mixture of 50 mM Tris pH 7.5 and 50 mM HEPES.

For GST or SFB pulldown analysis, cell lysates were prepared in pulldown lysis buffer including 150 mM NaCl, 1 mM Na_3_O_4_V, 10 mM NaF, 10 mM β-glycerolphosphate, 1% NP-40, 1 mM DTT, 1 mM PMSF, 1 μg/ml aprotinin, and 1 μg/ml leupeptin in 50 mM Tris pH 8.0. The cell lysates in this lysis buffer were treated with 50 units each of micrococcal nuclease (MNase, Worthington) and Benzonase nuclease (Merck) for ∼ 30 minutes at 37°C to maximize protein extraction, possibly including chromatin-bound proteins, and the enzyme activity was terminated by adding 5 mM EDTA. These samples were added to GST-beads incubated with overexpressed proteins or Streptavidin Sepharose beads (Cytivia). The bead-bound proteins, after washing with RIPA buffer containing 1% Triton X-100, 1% sodium deoxycholate, 0.1% SDS, 2 mM EDTA in PBS, were subjected to Western blot analysis.

To detect XRCC1 acetylation, a denaturing step was added to the pulldown Western blot (Suppl. Fig. 5A). The cell lysates in denaturing solution containing 50 mM Tris pH7.5, 1% SDS, 5 mM EDTA and 0.07% β-mercaptoethanol underwent a boiling and cooling cycle, followed by incubation in RIPA-based buffer containing 1% Triton X-100, 1% sodium deoxycholate, 0.1% SDS, 2 mM EDTA, 1 mM Na_3_O_4_V, 10 mM NaF, 10 mM β-glyerolphosphate, 1 mM DTT, 1 mM PMSF, 1 μg/ml aprotinin, 1 μg/ml leupeptin in PBS. The final supernatants were used for SFB pulldown followed by Western blot analysis.

Western blot samples were separated on SDS-PAGE gels, transferred onto nitrocellulose membranes (Cytiva) and visualized by enhanced chemiluminescence (ECL, Cytiva or Bio-Rad Laboratories) after incubation with indicated antibodies (Suppl. Table 3). The amounts of loaded protein were adjusted based on protein quantification by the Bradford method (Bio-Rad Laboratories), Ponceau S staining, Actin/Tubulin Western blot. To analyze DNA damage response (DDR), DNA damage was induced in targeted cells by treatment with reagents (at the dose and duration of each drug as indicated in the figures), and cells were collected immediately after drug treatment (0-hour time point), 3 hours, or when necessary 12 hours after removing drugs.

### Cell cycle analysis

The target genes were overexpressed or knocked down in cell lines by transfection of cloned gene constructs or siRNA primer pairs. Forty eight hours after transfection, cells were exposed to DNA damage inducing reagents as indicated in the figures, such as 2 hours for CPT or 24 hours for other reagents, and then fixed with 75% ethanol at −20°C 24 hours after initiation of DNA damage. On the following day, fixed cells, which were washed with PBS and treated with RNase A (100 μg/ml, Macherey-Nagel) were stained with propidium iodide (PI) for cell cycle analysis by Fluorescence Activated Cell Sorting (FACS, FACSCanto II, BD Biosciences). Ten thousand cells for each experimental group were measured and analyzed by FACSDiva software (v.9.2, BD Biosciences).

### Colony formation assay

Targeted cells were seeded at 3,000 cells per well in 6-well culture plates and maintained under the culture condition described in the ‘cell culture’ section above. DNA damage treatment was initiated 48 hours post-transfection and continuously refreshed with culture media changes for approximately 10 days. Afterward, the cell colonies were fixed with 100% methanol and stained with 0.01% Crystal Violet solution. Each well of the completely dried 6-well plates was photographed, and the area of stained colonies in each well image was subjected to measurements by ImageJ (v1.5, NIH, USA).

### Sulforhodamine B (SRB) assay

Twenty four hours after transfection, targeted cells were seeded at 500 cells per well in 96-well culture plates. Another 24 hours later, DNA damage treatment was applied, and the 96-well plates were fixed with 10% trichloroacetic acid (TCA, Sigma-Aldrich) solution on every alternate day for 6 days (days 2, 4, 6). Additionally, 96 well plates were immediately fixed after 1 hour of exposure to DNA damage (0-day time point). MMS, H_2_O_2_ and Phleo were refreshed once for cells with overexpression of target genes during the course of the experiment. CPT was removed 2 hours after treatment from siRNA-transfected cells. Fixed cells were stained with 0.4% Sulforhodamine B (SRB, Sigma-Aldrich) solution in 1% acetic acid for 30 minutes without light exposure. The SRB stained cells were dissolved in 10mM Trizma base (Sigma-Aldrich), and the color intensity was measured by a microplate spectrophotometer (BioTek plate reader, Agilent Technologies) at 540 nm with GEN5 data analysis software (v3.0, Agilent Technologies).

### Cell death assay

For cell death analysis *in vitro*, the Annexin V apoptosis detection method (ThermoFisher) was applied to DNA damaged cells according to the manufacturer’s guidelines. Approximately 2 × 10^6^ cells were exposed to DNA damaging condition as indicated in figure 3. All procedures were conducted in EDTA or calcium chelator free buffer. PI staining was added to detect loss of membrane integrity. Populations of Annexin V and PI positive cells were measured by FACS. Annexin V positive and PI negative cells were considered early apoptotic cells, while Annexin V and PI double positive cells were considered late apoptotic cells.

### Micro-irradiation (Micro-IR) analysis and Immunocytochemistry (ICC)

Cells transfected with GFP or mCherry-tagged gene constructs were seeded onto glass bottom cell culture dishes (SPL Life Sciences). The cell numbers were adjusted to reach ∼80% confluency one day before micro-irradiation (micro-IR) when BrdU (10 μM, Sigma-Aldrich) was added to the culture. To knock down target genes and identify targeted cells, siRNA primers labelled with 5’-cyanine 3 (Cy3) were used to transfect cells 48 hours prior to micro-IR. The condition conducted with a 405 nm laser (Nikon) was optimized for XRCC1 recruitment to the micro-IR site faster than the known DNA DSB responder RSF1 (data not shown) ^40^. Accumulation of fluorescent signals at the micro-IR sites was observed under a spectral confocal laser dual scanning microscope (Eclipse A1Rsi and Eclipse Ti-E, Nikon) equipped with an environmentally controlled chamber (temperature, humidity, and CO_2_) installed at the 3D immune system imaging core facility, Ajou University Medical Center. The real-time accumulation and intensity kinetics of GFP signals at DNA damage site were tracked for 3 minutes and analyzed using NIS-elements software (v5.21, Nikon).

GCN5 and TADA2B were not feasible for micro-IR analysis in real-time. Therefore, GFP or mCherry-tagged GCN5 or TADA2B accumulation at the DNA damage sites were examined by immunocytochemistry (ICC). Targeted cells were fixed with 2% formaldehyde solution (Sigma-Aldrich) 10 minutes after micro-IR, and GFP/mCherry localization was imaged with the same microscopic equipment. γH2AX immunoreactivity visualized by Cy3, FITC, or Alexa Fluor 647 secondary antibody was used as an indicator of DNA damage. Furthermore, the retention of XRCC1 at the micro-IR site was examined 1 and 15 minutes after DNA damage induction via ICC.

In addition, γH2AX foci formation after DNA damage induction by drug treatment was analyzed to estimate damage repair. Transfected cells were seeded onto round cover glasses (Paul Marienfeld), and DNA damaged cells were fixed in 1% PBS-buffered paraformaldehyde 30 minutes (10 minutes in the case of H_2_O_2_), 6 hours, and 24 hours after initiation of drug treatment as indicated in the figures. γH2AX foci were visualized by Cy3 secondary antibody and the nuclei were stained with 4’,6-diamidino-2-phenylindole (DAPI). Microscopic images were examined and captured with a B600TiFL microscope (Optika) and a DFC130 digital camera (Leica Microsystems). The γH2AX foci formation was calculated as the ratio of γH2AX immunopositivity to the area of the nucleus measured using ‘QuPath’ (v0.5.1) ^41^ and analyzed further for statistical differences.

### DNA double strand break repair (DSBR) assay

Specific vector constructs were purchased to measure DSBR capacity. pHPRT-DRGFP (Addgene) was used to monitor homologous recombination repair (HRR), and pimEJ5GFP (Addgene) was used for non-homologous end joining (NHEJ). The pCBAScel (Addgene), which expresses I-SceI endonuclease, was used to introduce a DSB at an I-SceI site located within the GFP site of the pHPRT-DRGFP or at two I-Sce1 sites flanking pgkPURO cassette which separates the pCAGGS promoter and eGFP in the pimEJ5GFP construct.

U2OS cells were transfected with *XRCC1*, *GCN5,* or *TADA2B* siRNA using RNAiMax (Invitrogen). Simultaneously, either pHPRT-DRGFP or pimEJ5GFP construct was transfected using PEI (Polysciences). Twenty four hours later, additional expression of pCBASceI using lipofectamine 2000 (Invitrogen) in Opti-MEM (Gibco), as instructed in the user’s manual, was introduced on the U2OS cells. GFP-positive cells, which indicate successful DSBR, were quantified by flow cytometry 2 days after the final transfection.

### Statistical analysis

All statistical analyses were performed using Prism software (v6.0, GraphPad). All numerical data were subjected to statistical analysis such as ANOVA followed by multiple comparisons or t-test with Welch’s corrections. P values less than 0.05 were considered significant (* p<0.05, ** p<0.01, *** p<0.005, **** p<0.001).

## RESULTS

### TADA2B and GCN5 are novel XRCC1 interacting proteins

To identify XRCC1 binding proteins, we first tested the feasibility of a GST-XRCC1 pulldown assay using mouse embryonic brain lysates. Well-known XRCC1 binding partners LIG3, PNKP, and POLB ^18^ were detected in eluates from glutathione columns saturated with GST-fused mouse XRCC1 amplified in insect cell culture (Fig. 1A). We then applied this GST pulldown assay to cerebellar lysates of adult *Xrcc1^WT^* and *Xrcc1^Nes-Cre^*mice brains. Compared to the GST alone and *Xrcc1^Nes-Cre^* samples, there were multiple unique bands detected in GST-XRCC1 pulldown samples. Mass spectrometry analysis identified some known interactors to XRCC1 (data not shown), and one unique band that was identified as TADA2B (Fig. 1A). This protein binds to GCN5 as part of the HAT module in the SAGA complex ^24,42^. TADA2B showed higher expression in the cerebellum than in the cerebral cortex, similar to XRCC1 and LIG3, while GCN5 exhibited the reverse pattern in the mouse brain. The expression of these two proteins was not affected by *Xrcc1* deficiency in the mouse brain (Fig. 1B). To facilitate detailed mechanistic analysis and to determine whether these novel XRCC1 interactions have relevance for DNA repair disorders, we proceeded to characterize these interactions using human proteins.

**Figure 1.**
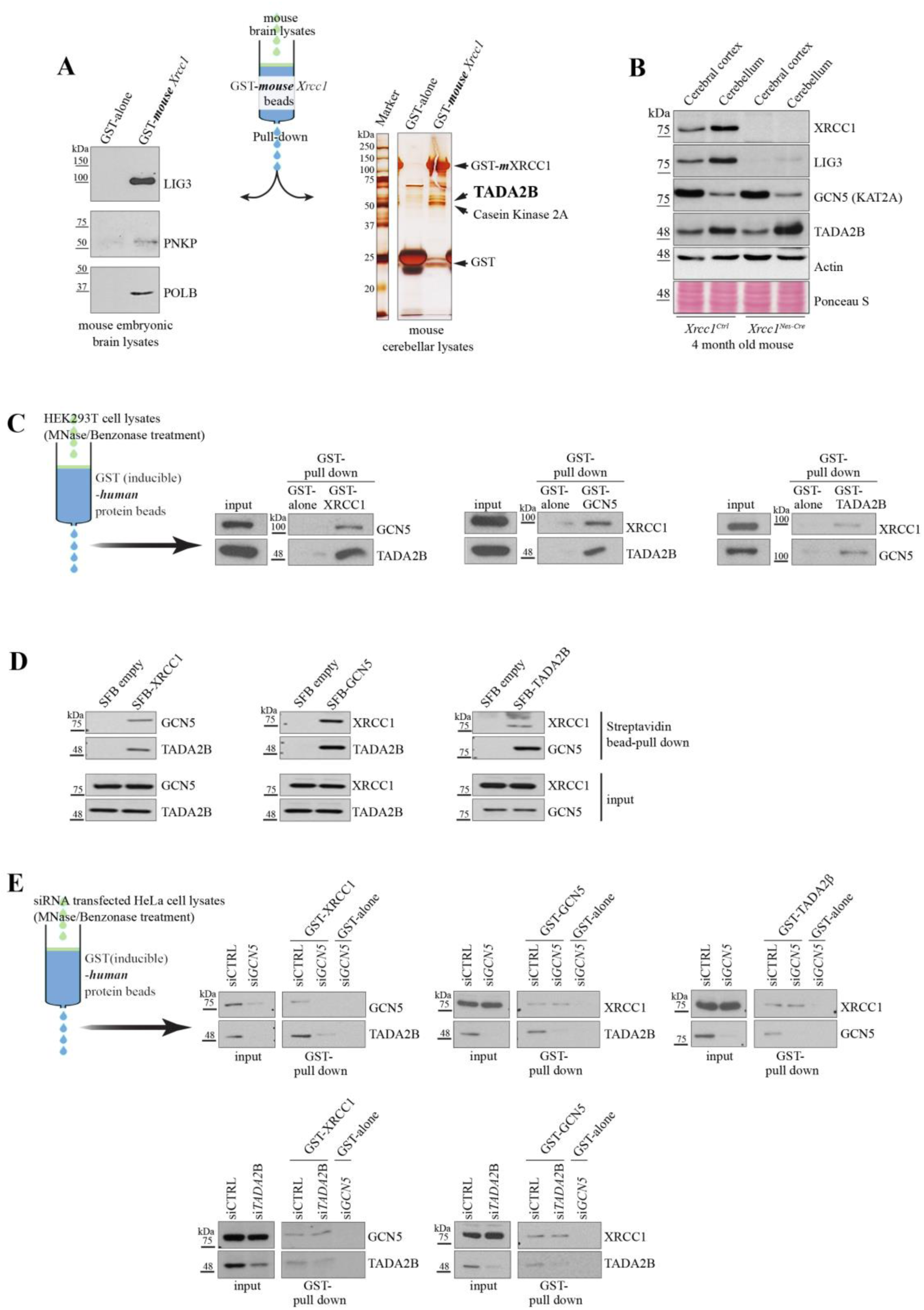
GCN5 and TADA2B are newly identified as novel XRCC1 interacting proteins. **A.** GST pulldown assays demonstrate XRCC1 binding partners in mouse brain tissue. Left panel: Western blot analysis confirms known XRCC1-interacting proteins (LIG3, PNKP, POLB) are captured by GST-mouse XRCC1 from embryonic brain lysates. Right panel: Silver staining of proteins pulled down from adult mouse cerebellar lysates reveals additional binding partners. Proteins identified by mass spectrometry are indicated, including TADA2B. **B.** Regional expression analysis of GCN5 and TADA2B in mouse brain. Western blot analysis shows normal expression of GCN5 and TADA2B proteins in *Xrcc1* null cerebral cortex and cerebellum, with distinct expression patterns between brain regions. Note the differential distribution: TADA2B, LIG3, and XRCC1 show higher expression in cerebellum, while GCN5 exhibits the opposite pattern. **C.** Validation of human protein interactions using cell-free GST pulldown assays. GST-human XRCC1 immobilized on glutathione beads captures endogenous GCN5 and TADA2B from nuclease-treated HEK293T cell lysates (MNase and Benzonase treatment to remove chromatin). Western blot analysis confirms reciprocal interactions among all three proteins. **D.** Confirmation of protein interactions in cellular context using streptavidin pulldown assays by Western blot. SFB-tagged XRCC1 overexpressed in HEK293T cells captures endogenous GCN5 and TADA2B, demonstrating these interactions occur within intact cells. **E.** Direct binding analysis using selective protein depletion by Western blot. GST-XRCC1 pulldown assays using HeLa cell lysates with targeted depletion of *GCN5* or *TADA2B* (siRNA knockdown) show XRCC1 binds directly to both proteins, as interactions persist in the absence of the third partner.

To confirm the interaction between XRCC1 and TADA2B, we implemented two complementary methods. First, GST-tagged human XRCC1, TADA2B, and GCN5 proteins were induced (Suppl. Fig. 1A) and passed over glutathione resin columns. MNase and Benzonase treated cell lysates were then applied to the columns to isolate XRCC1 binding proteins, allowing us to detect protein-protein interactions in a cell-free system. Initially, GCN5 was included as a positive control for protein-protein interaction with TADA2B. Unexpectedly, we observed that all three proteins, XRCC1, TADA2B, and GCN5 interact with each other (Fig. 1C). To validate these interactions within cells, we expressed SFB-tagged XRCC1, TADA2B, and GCN5 proteins (Suppl. Fig. 1B). Using streptavidin bead pulldown assays, we demonstrated that endogenous XRCC1, TADA2B, or GCN5 bound to SFB-tagged counterparts, consistent with the GST pulldown data and confirming that XRCC1 binds to both TADA2B and GCN5 (Fig. 1D).

Given the possble direct interaction between GCN5 and TADA2B, we investigated whether either protein might indirectly bind to XRCC1 through the other. We performed additional GST-pulldown assays using lysates from HeLa cells with either *GCN5* or *TADA2B* knockdown. siRNA mediated gene knockdown effectively reduced target protein levels (Suppl. Fig. 1C). Interestingly, *GCN5* knockdown also reduced TADA2B expression, which was restored by treatment with the proteasome inhibitor MG132, suggesting that TADA2B stability depends on GCN5 binding (Suppl. Fig. 1D), reminiscent of the XRCC1-LIG3 relationship (Fig. 1B and Suppl. Fig. 11A and 11D). When *GCN5* was knocked down, neither GCN5 nor TADA2B was pulled down by GST-XRCC1. However, both GST-GCN5 and GST-TADA2B successfully pulled down endogenous XRCC1 in *GCN5* knockdown cell lysates, which lack endogenous TADA2B due to its destabilization. Moreover, XRCC1-GCN5 direct binding was detected in *TADA2B* knockdown samples (Fig. 1E). These data indicate that XRCC1 interacts directly with both GCN5 and TADA2B.

### The BRCT domains of XRCC1 serve as the primary binding sites for GCN5 and TADA2B

We further characterized the specific binding domains in these three proteins. Based on previous reports ^43–45^, we cloned and generated several truncated variants of each protein (Suppl. Fig. 2A to 2C). We also created XRCC1 proteins lacking either the BRCT I domain (XRCC1 ΔB1D) or BRCT II domain (XRCC1 ΔB2D), given the known importance of BRCT domains in protein-protein interactions ^18,46,47^ (Suppl. Fig. 2D). Streptavidin pulldown assays demonstrated that GCN5 could bind to the XRCC1 NTD and BRCT I domains, while TADA2B interacts with the XRCC1 NTD and both BRCT domains (Fig. 2A). The binding of GCN5 and TADA2B to XRCC1 BRCT domains was also confirmed in cell-free GST-XRCC1 pulldown assay (Suppl. Fig. 3A). To further validate these domain specific interactions, particularly the BRCT domains, we performed streptavidin pulldown using SFB-XRCC1 ΔB1D or SFB-XRCC1 ΔB2D. These experiments clearly showed that the XRCC1 BRCT I domain is the primary binding site for GCN5, whereas the XRCC1 BRCT II domain specifically binds to TADA2B (Fig. 2B and Fig. 6A). Reciprocal domain mapping for protein interactions revealed that XRCC1 interacts primarily with the GCN5 ACT (HAT functional domain) and the TADA2B CTL (Suppl. Fig. 3B and 3C). The interaction sites between GCN5 and TADA2B were consistent with previous reports, though additional binding interfaces were detected ^44,48^. To complement our wet-laboratory data, we employed web-based structure prediction tools (AlphaFold and ColabFold) to model potential interactions of three proteins ^49,50^. One prediction showed GCN5 in close proximity to the BRCT I domain and TADA2B adopting an extended conformation spanning both BRCT I and II domains (Suppl. Fig. 3D). These computational predictions corroborate our experimental findings, which demonstrate distinct binding preferences of the two BRCT domains of XRCC1 for GCN5 and TADA2B.

**Figure 2.**
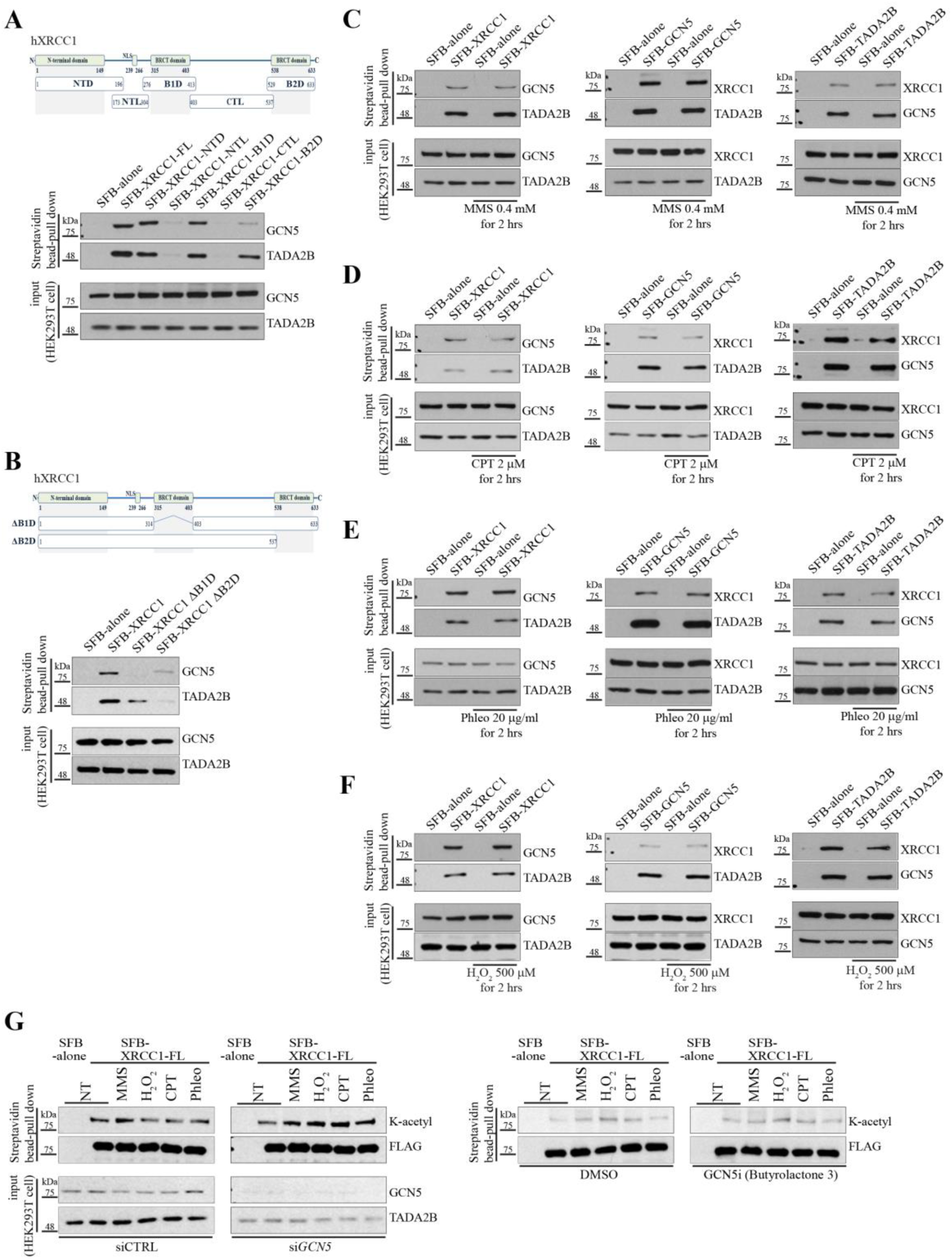
The binding of GCN5 or TADA2B to XRCC1 is constitutively maintained and does not induce XRCC1 modification. **A.** Domain mapping of XRCC1-GCN5/TADA2B interactions. Upper panel: Schematic diagram of truncated *XRCC1* constructs showing N-terminal domain (NTD), BRCT I domain (B1D), and BRCT II domain (B2D). Lower panel: Western blots of streptavidin pulldown assays with SFB-tagged truncated XRCC1 proteins demonstrate that GCN5 interacts with both NTD and BRCT I domains, while TADA2B binds to NTD and both BRCT domains. **B.** BRCT domain specific binding analysis. Upper panel: Schematic diagram of XRCC1 deletion mutants lacking either BRCT I (ΔB1D) or BRCT II (ΔB2D) domains. Lower panel: Western blots of streptavidin pulldown experiments reveal that GCN5 binding requires the BRCT I domain, while TADA2B binding depends primarily on the BRCT II domain. **C∼F.** DNA damage does not alter protein complex stability. HEK293T cells expressing SFB-tagged proteins were treated with DNA damaging agents; methyl methanesulfonate (MMS) (C), camptothecin (CPT) (D), phleomycin (Phleo) (E), or hydrogen peroxide (H_2_O_2_) (F) at indicated concentrations. Western blots of streptavidin pulldown assays immediately after 2 hour exposure to DNA damaging drugs demonstrate that GCN5, TADA2B, and XRCC1 interactions remain unchanged following DNA damage induction, indicating constitutive complex formation. **G.** GCN5 does not acetylate XRCC1 in response to DNA damage. Left panel: Analysis of XRCC1 lysine (K-acetyl) acetylation following DNA damage treatment (MMS, H_2_O_2_, CPT, or Phleo at the same dose and exposure time as in figure 2C∼2F) shows no induction of acetylation. siRNA mediated *GCN5* depletion (which also reduces TADA2B levels due to protein instability) does not affect XRCC1 acetylation status upon DNA damage. Right panel: Pharmacological inhibition of GCN5 acetyltransferase activity (butyrolactone 3) similarly fails to alter XRCC1 acetylation levels, confirming that XRCC1 is not a direct substrate of GCN5.

### DNA damage neither enhances protein interactions nor induces GCN5-mediated XRCC1 acetylation

Typically, XRCC1 binding partners show enhanced interaction with XRCC1 following DNA damage ^18,19^. Therefore, we tested whether GCN5 or TADA2B binding to XRCC1 is modulated by DNA damage. We employed four different DNA damaging reagents: Methyl methanesulfonate (MMS – alkylating agent), Camptothecin (CPT – topoisomerase I inhibitor inducing strand break damage), Phleomycin (Phleo – a radiomimetic drug), and Hydrogen peroxide (H_2_O_2_ – oxidative stress causing strand break damage), each inducing distinct types of DNA lesions ^51^. None of these drug treatments affected protein interaction among XRCC1, GCN5, and TADA2B. Both U2OS and HeLa cells overexpressing SFB-tagged proteins were subjected to pulldown assay before and after DNA damage induction. All three proteins were pulled down together under normal conditions without stress, and these protein interactions remained unchanged following exposure to DNA damaging agents (Fig. 2C-2F and Suppl. Fig. 4A-4D). These findings suggest that XRCC1 constitutively binds to GCN5 and TADA2B regardless of DNA damage status.

Having established the constitutive nature of these protein interactions, we next investigated whether XRCC1 serves as a substrate for GCN5 HAT enzyme activity ^24^. To assess potential changes in XRCC1 acetylation before and after DNA damage, we implemented a denaturing protocol to remove any XRCC1 associated proteins prior to streptavidin pulldown and Western blot detection with anti-acetyl lysine (K) antibody (Suppl. Fig. 5A). Our results showed no significant induction of XRCC1 acetylation following DNA damage. Moreover, inhibition of GCN5 activity, either pharmacologically (GCN5 inhibitor MB-3/Butyrolactone 3) or through siRNA mediated gene knockdown, did not alter the status of XRCC1 acetylation (Fig. 2G).

Collectively, these data demonstrate that the interactions among XRCC1, GCN5, and TADA2B deviate from conventional paradigms in DNA damage repair mechanisms. Unlike typical repair protein interactions that are enhanced by DNA damage, these interactions remain constitutive. Furthermore, despite GCN5’s acetyltransferase function, XRCC1 does not appear to be a direct substrate, even though XRCC1 binds to the HAT containing domain of GCN5 (Suppl. Fig. 3B).

### Depletion of GCN5 or TADA2B sensitizes cells to DNA damage

To investigate the functional roles of GCN5 and TADA2B in DNA repair and responses, we analyzed the consequence of DNA damage under two conditions: Gain- or Loss-of-function conditions. We examined long-term effects of DNA damage on proliferation using colony forming and SRB assays. Knockdown of *XRCC1*, *GCN5*, or *TADA2B* individually slowed down overall proliferation compared to controls (Suppl. Fig. 6D), whereas overexpression of these genes did not affect proliferation rates (data not shown). In colony forming assays, U2OS cells with gene knockdown showed enhanced sensitivity to DNA damaging agents in a dose dependent manner compared to controls. *TADA2B* knockdown cells showed reduced proliferation following phleomycin or H_2_O_2_ treatment, while *XRCC1* knockdown cells displayed higher sensitivity to MMS exposure compared to other knockdown cell lines (Fig. 3A). This heightened sensitivity to different DNA damaging reagents was not observed in cells overexpressing these three genes (Suppl. Fig. 6A). Similarly, SRB assays conducted with the indicated dose of DNA damaging reagents revealed significantly reduced survival in all three knockdown cell lines relative to control groups (Fig. 3B), these effects were not detected in cells overexpressing these proteins (Suppl. Fig. 6B). These data demonstrate that reduction of GCN5 or TADA2B renders cells vulnerable to DNA damage, suggesting roles for GCN5 and TADA2B in DNA damage repair or responses.

**Figure 3.**
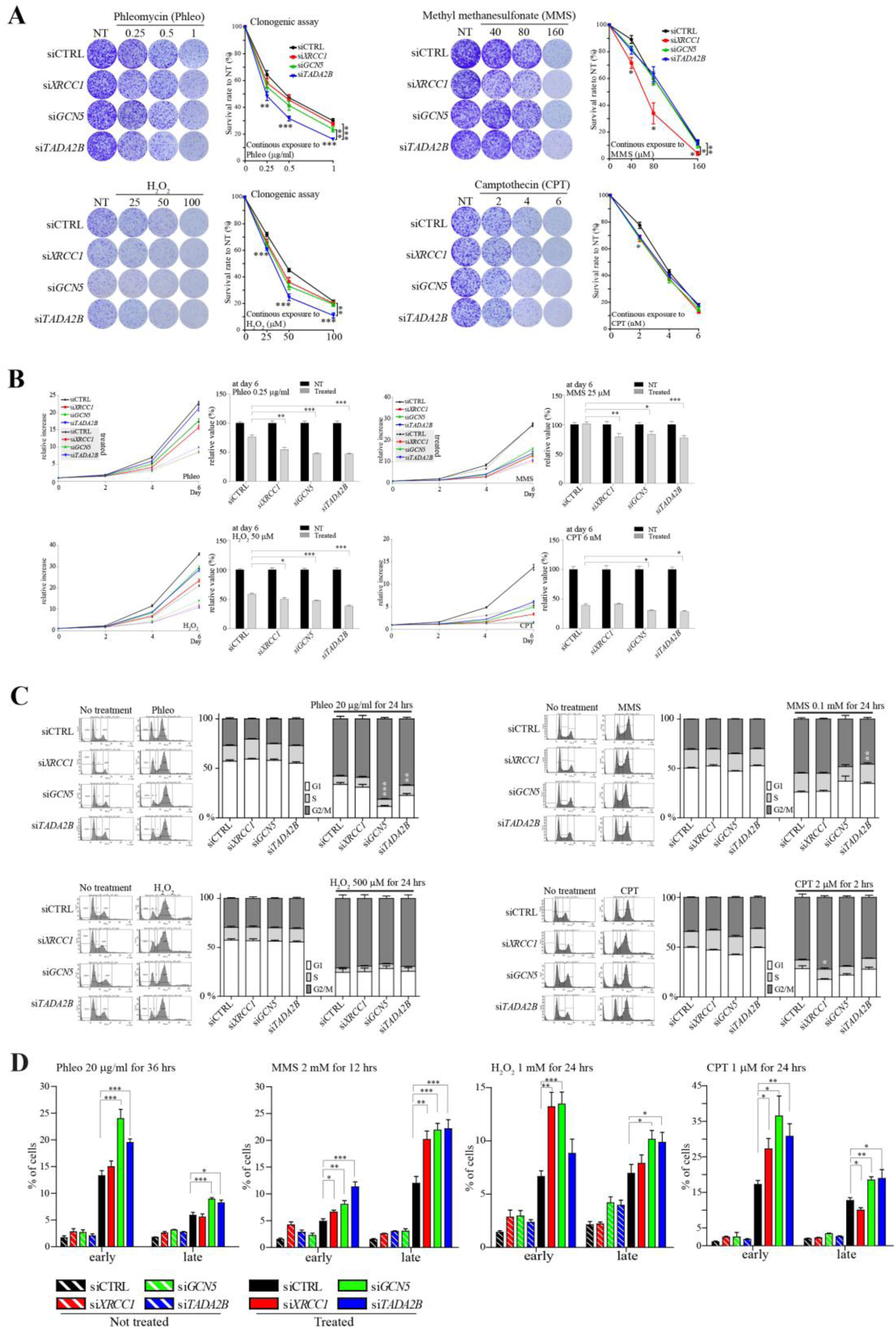
*GCN5* or *TADA2B* depletion reduces cell viability. **A.** Colony formation assay demonstrates reduced survival following DNA damage. Control (CTRL) and gene depleted U2OS cells (*XRCC1*, *GCN5*, or *TADA2B* siRNA) were continuously exposed to increasing concentrations of phleomycin (Phleo), methyl methanesulfonate (MMS), hydrogen peroxide (H_2_O_2_), or camptothecin (CPT) for 9-10 days. Left panels: Representative Crystal Violet staining shows reduced colony formation in gene depleted cells. Line graphs in right panels: Quantification reveals dose-dependent survival defects in all three knockdown lines, with distinct sensitivity profiles to different DNA damaging agents. N=6; ***: p<0.005, **: p<0.01, *: p<0.05. **B.** Sulforhodamine B (SRB) assay confirms growth defects following DNA damage. Control and gene depleted U2OS cells (*XRCC1*, *GCN5*, or *TADA2B* siRNA) were treated with DNA damaging agents, phleomycin (Phleo), methyl methanesulfonate (MMS), hydrogen peroxide (H_2_O_2_), or camptothecin (CPT), at indicated doses and time points. Line graphs in left panels: Growth curves show impaired proliferation in knockdown cells relative to day 0. Bar graphs in right panels: Relative viability at day 6 demonstrates significant reduction in survival for all depleted cell lines compared to untreated controls. N=6; ***: p<0.005, **: p<0.01, *: p<0.05. **C.** Cell cycle analysis reveals normal checkpoint activation despite increased sensitivity. FACS analysis of control and gene depleted U2OS cells (*XRCC1*, *GCN5*, or *TADA2B* siRNA) following 24 hour exposure to DNA damaging agents at indicated concentrations. Histograms in left panels: Representative cell cycle profiles from FACS analyses. Bar graphs in right panels: Quantification of cell cycle distribution (G1, S and G2/M phases) before and after exposing to DNA damaging reagents. N=6; ***: p<0.005, **: p<0.01, *: p<0.05 **D.** Cell death analysis shows either GCN5 or TADA2B knockdown sensitize cells to DNA damage. Histograms show percentages of Annexin V/PI positive cells by FACS analysis. Early stage apoptotic cells are Annexin V positive and PI negative populations, and late stage apoptotic cells are Annexin V/PI double positive populations. Control and gene knockdown U2OS cells (*XRCC1*, *GCN5*, or *TADA2B* siRNA) were exposed to DNA damaging conditions as indicated in the figure. N=4∼6; ***: p<0.005, **: p<0.01, *: p<0.05

Cell cycle arrest is one of the consequences resulting from DNA damage. Therefore, we examined potential alterations in cell cycle progression in three knockdown cell lines. When exposed to DNA damaging reagents, all cells displayed G2/M arrest (Fig. 3C). *XRCC1* knockdown cells exhibited a similar arrest levels to controls, except following CPT treatment, to which these cells show known heightened sensitivity ^20^. *GCN5* and *TADA2B* knockdown cells showed enhanced G2/M accumulation compared to controls when exposed to the strand break inducer Phleo. Overall, knockdown of *XRCC1*, *GCN5*, or *TADA2B* did not significantly disrupt the cell cycle arrest response to DNA damage compared to control cells (Fig. 3C). And no differences in G2/M arrest following DNA damage induction were found in cells overexpressing these proteins (Suppl. Fig. 6C). DNA damage ultimately triggers apoptotic cell death when repair mechanisms are overwhelmed. To assess this endpoint, we quantified apoptotic cell populations using Annexin V/PI staining and FACS analysis. Following exposure to various DNA damaging agents, all three gene knockdown cells exhibited elevated populations of early and late stage apoptotic cells (Fig. 3D and Suppl. Fig. 7), implying GCN5 and TADA2B knockdown compromises cellular DNA damage tolerance, sensitizing cells to genotoxic stress.

### Inactivation of GCN5 or TADA2B impairs DNA damage repair efficacy

Since both *GCN5* and *TADA2B* knockdown rendered cells vulnerable to DNA damage, we looked into the early signaling events upon DNA damage induction and damage repair efficacy. All three knockdown cell lines showed time course kinetics of DDR events comparable to control cells, including ATM phosphorylation and phosphorylation of its kinase substrates such as KAP1, CHK2, p53, and H2AX, as well as CHK1 phosphorylation (Fig. 4A to 4D). These DDR kinetics remained also unaltered in cells overexpressing these genes (Suppl. Fig. 8A to 8D). However, all three knockdown cells maintained elevated γ-H2AX (phosphorylated H2AX) levels 3 hours after removal of the DNA damage agents, suggesting impaired DNA repair efficiency and implying roles for GCN5 and TADA2B in DNA damage repair.

**Figure 4.**
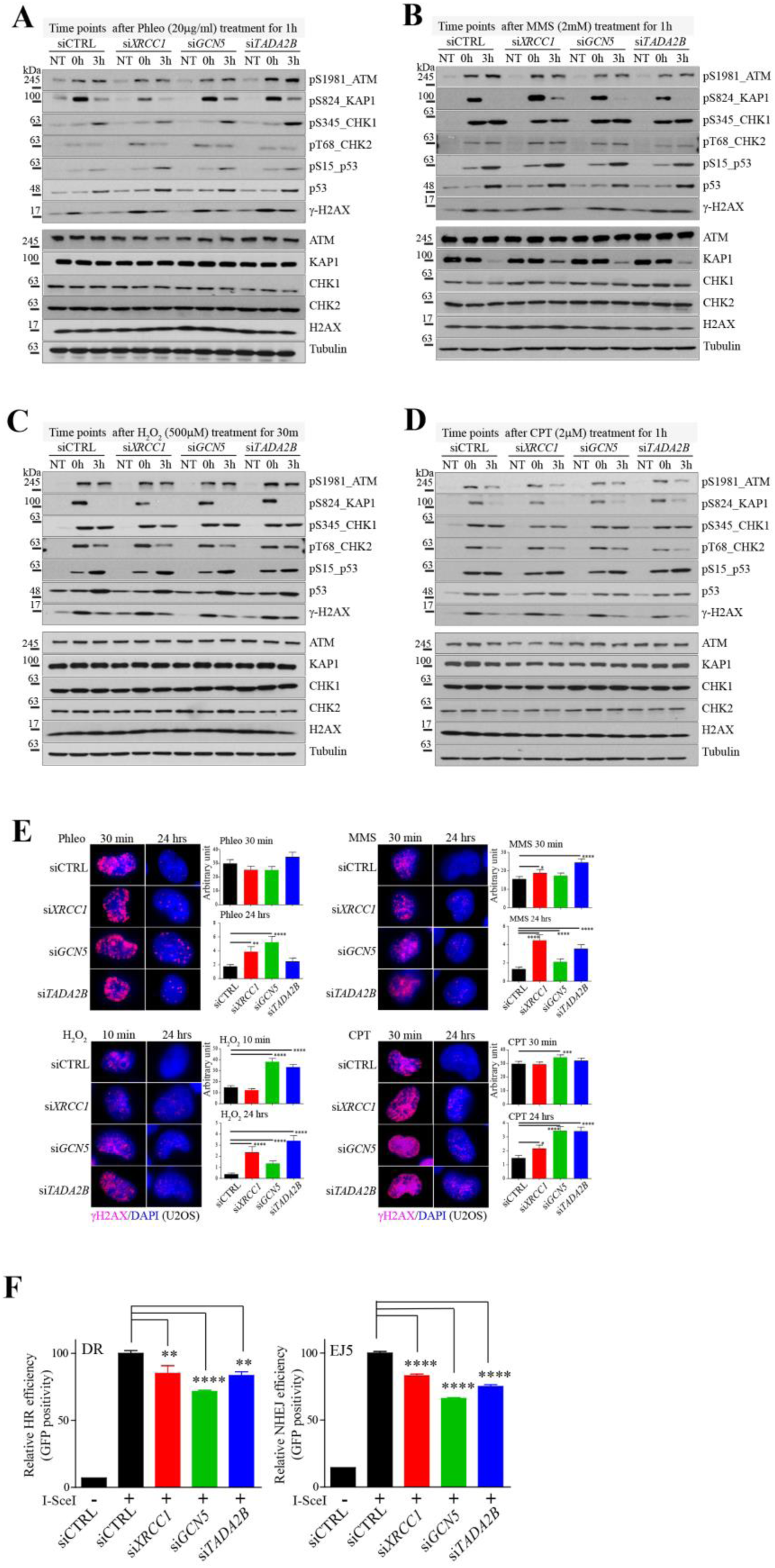
GCN5 or TADA2B depletion results in defective DNA damage repair. **A∼D.** DNA damage response signaling remains intact in gene depleted cells. Western blot analysis of control (CTRL) and gene depleted U2OS cells (*XRCC1*, *GCN5*, or *TADA2B* knockdown) following DNA damage induction. Cells were treated with phleomycin (A; Phleo, 20 μg/ml, 1 hour), methyl methanesulfonate (B; MMS, 2 mM, 1 hour), hydrogen peroxide (C; H_2_O_2_, 500 μM, 30 minutes), or camptothecin (D; CPT, 2 μM, 1 hour) and collected immediately (0h) or 3 hours post-treatment (3h). NT: no treatment. **E.** Persistent DNA damage foci demonstrate defective repair kinetics. Immunocytochemistry analysis of γ-H2AX foci formation and resolution following brief DNA damage pulses: phleomycin (Phleo, 20 μg/ml, 30 min), MMS (1 mM, 30 min), H_2_O_2_ (500 μM, 10 min), or camptothecin (CPT, 1 μM, 30 min). Left panels: Representative images show γ-H2AX foci at indicated time points post-damage. Bar graphs in right panels: Quantitative analysis at immediately after removing DNA damaging reagents and 24 hours post-treatment reveals significantly delayed foci resolution in all three gene-depleted U2OS cell lines (*XRCC1*, *GCN5*, or *TADA2B* siRNA), implying impaired DNA repair capacity. N>300 nuclei per condition; ****: p<0.001, ***: p<0.005, **: p<0.01, *: p<0.05. **F.** DNA strand break repair pathways are compromised in gene depleted cells. Functional repair assays using fluorescent reporter constructs in control (CTRL) and gene depleted U2OS cells (*XRCC1*, *GCN5*, or *TADA2B* siRNA). Cells were transfected with pHPRT-DRGFP (DR, homologous recombination reporter) or pimEJ5GFP (EJ5, non-homologous end joining reporter) constructs, and repair efficiency was measured by GFP-positive cell populations following I-SceI-induced DNA breaks. All three knockdown cell lines show significant defects in both DNA strand break repair pathways. N=6; ****: p<0.001, **: p<0.01.

During these analyses, we observed unexpected regulation of KAP1 (TRIM28), a protein that involved in chromatin decondensation near DNA break sites ^52^. Total KAP1 protein levels decreased progressively following MMS treatments specifically, without affecting its phosphorylation status (Fig. 4B and Suppl. Fig. 8B), similar to a previous report ^53^. Interestingly, this reduction was attenuated in *XRCC1* knockdown cells (Fig. 4B), while XRCC1 overexpression enhanced KAP1 phosphorylation even without DNA damage, and KAP1 phosphorylation levels after exposure to all four DNA damaging reagents were enhanced compared to other cell lines (Suppl. Fig. 8A to 8D). These data suggest that XRCC1 may play a novel function in chromatin regulation. While this KAP1 regulation by XRCC1 is beyond the scope of the current study, it may warrant future investigation.

To further validate these repair defects, we monitored changes in γ-H2AX by ICC over 24 hours after removal of DNA damaging reagents (Fig. 4E and Suppl. Fig. 9A to 9D). Following DNA damage induction, *XRCC1* and *TADA2B* knockdown cells exhibited heightened sensitivity to MMS treatment. Both *GCN5* and *TADA2B* knockdown cells were significantly more susceptible to H_2_O_2_ treatment than the other cell lines. At 24 hours post-treatment, all three knockdown cell lines failed to resolve γ-H2AX foci, demonstrating persistent unrepaired DNA lesions and establishing that both GCN5 and TADA2B are functionally essential for efficient DNA repair. Furthermore, we measured the capacity to repair DR-GFP or EJ5-GFP constructs with strand breaks induced by I-SceI restriction enzyme in the three knockdown cell lines (Fig. 4F). All three gene knockdown cell lines showed impaired HRR and NHEJ repair efficiency. While XRCC1 is not directly involved in DSBR, unrepaired SSB can be converted to DSBs during replication or through processing errors ^18,19^. Therefore, the defective repair capacity observed in these assays reflects broader abnormalities in DNA repair mechanisms. These data provide compelling evidence that GCN5 and TADA2B are not merely XRCC1 binding partners, but critical regulators whose loss compromises DNA repair capacity.

### GCN5 and TADA2B control XRCC1 recruitment and retention at DNA damage sites

Next, we examined how XRCC1 behavior is affected by GCN5 and TADA2B. We applied a technique that allows real-time tracking of protein movement after DNA damage induced by micro-IR. For this application, XRCC1, GCN5, and TADA2B were tagged with fluorescent markers (GFP or mCherry) (Suppl. Fig. 5B and 5C). In addition, XRCC1 ΔB1D and ΔB2D were fluorescently tagged (Suppl. Fig. 5D). Although we could not detect recruitment of GCN5 and TADA2B to micro-IR sites in real-time, ICC revealed that both GCN5 and TADA2B were localized at DNA damage sites after micro-IR, suggesting that GCN5 and TADA2B respond to DNA damage and are likely involved in DNA damage repair processes (Fig. 5A). We then examined XRCC1 recruitment in real-time after micro-IR. Fast and focused accumulation of XRCC1 at micro-IR sites was significantly attenuated in either GCN5 or TADA2B knockdown conditions (Fig. 5B), suggesting that constitutive protein interactions between GCN5/TADA2B and XRCC1 are required for proper XRCC1 response to DNA damage. Furthermore, loss of the BRCT I domain completely prevented XRCC1 recruitment, while XRCC1 lacking the BRCT II domain showed impaired recruitment to micro-IR sites (Fig. 5C). These recruitment defects of XRCC1 BRCT domain mutants persisted in either GCN5 or TADA2B knockdown cells (Fig. 5D and 5E). We also analyzed whether GCN5 enzyme activity inhibition could influence XRCC1 recruitment. Butyrolactone 3 treatment did not change the course of XRCC1 and XRCC1 BRCT domain mutant recruitments (Suppl. Fig. 10A and 10B), consistent with evidence that GCN5 HAT activity is not involved in XRCC1 function.

**Figure 5.**
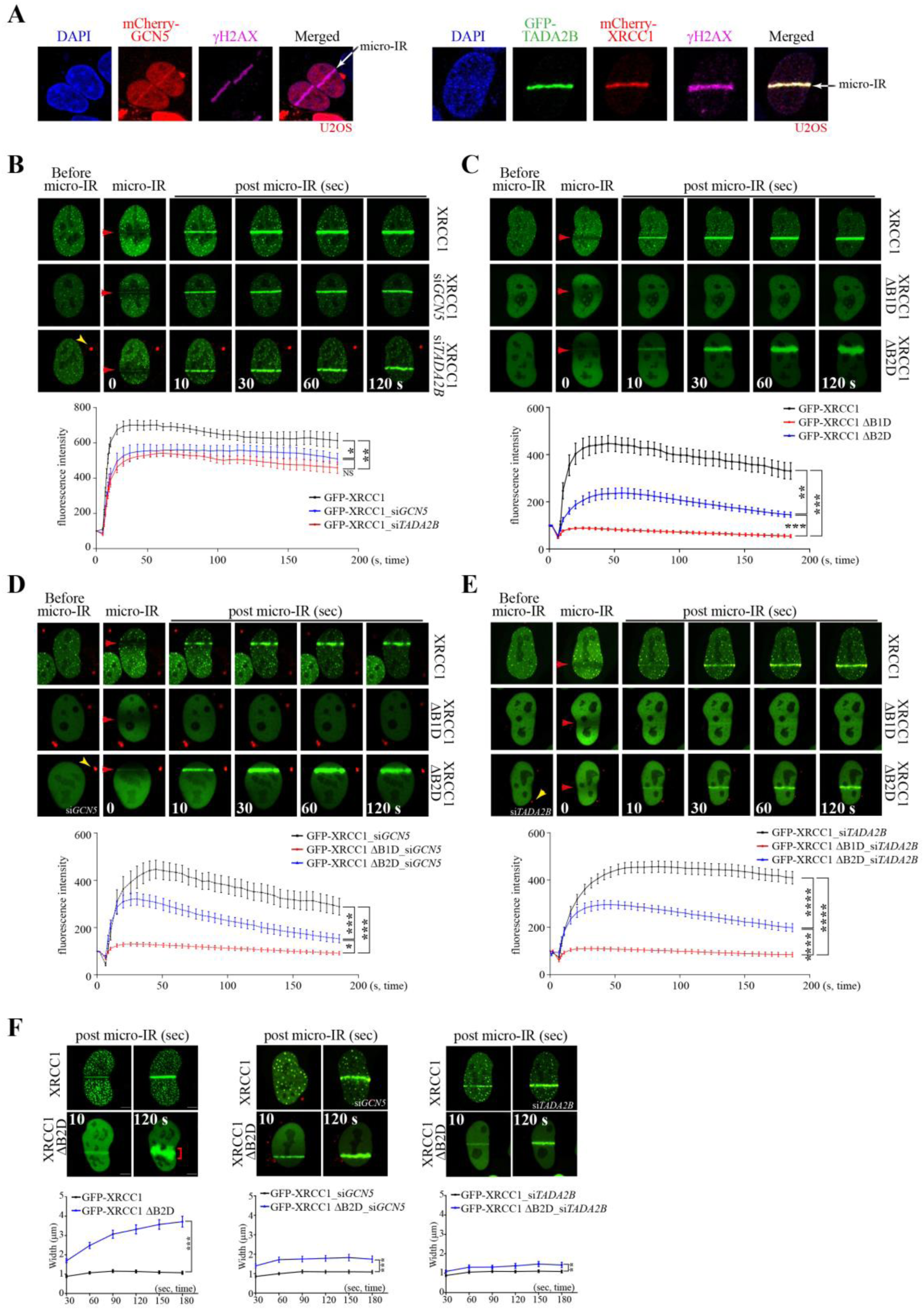
Recruitment of XRCC1 to DNA damage sites is defective in absence of GCN5 or TADA2B. **A.** GCN5 and TADA2B are recruited to DNA damage sites alongside XRCC1. Immunocytochemistry of U2OS cells following localized micro-IR shows recruitment of tagged GCN5, TADA2B, and XRCC1 proteins (GFP or mCherry) to damage sites by induced by micro-IR. γH2AX immunostaining (Alexa Fluor 647) marks micro-IR regions. All three proteins (GCN5, TADA2B, and XRCC1) accumulate at DNA damage sites, demonstrating their coordinated response to genotoxic stress. **B∼E.** Real-time analysis reveals impaired XRCC1 recruitment kinetics in *GCN5* or *TADA2B* depleted cells. Live cell imaging of GFP-tagged XRCC1 (WT and BRCT domain deletion mutants) following micro-IR in control and gene depleted U2OS cells (*GCN5* or *TADA2B* knockdown). Upper panels: Representative time course images showing GFP accumulation at damage sites (red arrowheads) over 180 seconds (s). Line graphs in lower panels: Quantitative kinetic analysis of recruitment efficiency. **B:** *GCN5* or *TADA2B* knockdown significantly impairs WT XRCC1 recruitment. **C:** BRCT I domain deletion (ΔB1D) abolishes XRCC1 recruitment completely, while BRCT II domain deletion (ΔB2D) reduce recruitment efficiency. **D-E:** defective recruitments of BRCT deletion mutants (ΔB1D and ΔB2D) maintain in *GCN5* or *TADA2B* depleted cells. Knockdown cells were identified by Cy3-labeled siRNA (yellow arrowheads); only Cy3-positive cells were analyzed. Most Cy3 positivity disappear during live cell imaging. N=8-10 cells per condition; ****: p<0.001, ***: p<0.005, **: p<0.01, *: p<0.05, NS: not significant. **F:** BRCT II domain is essential for stable XRCC1 focal accumulation and retention at damage sites and is regulate by GCN5/TADA2B. XRCC1 lacking the BRCT II domain (ΔB2D) shows progressive spreading of GFP signal at damage sites which is not observed in WT XRCC1 (left panel, ΔB2D slope = 0.01, WT slope = 0.001), indicating defective retention. This spreading phenotype is rescued in *GCN5* (middle panel, ΔB2D slope = 0.002, WT slope = 0.001) or *TADA2B* (right panel, ΔB2D slope = 0.002, WT slope = 0.001) depletion. Upper panels: Representative images showing GFP signal width (red bracket) over time. Line graphs in lower panels: Quantification of signal spreading demonstrates the retention defect and its rescue upon *GCN5/TADA2B* depletion. N=8-10 cells; ***: p<0.005, **: p<0.01.

An important finding was that XRCC1 ΔB2D was poorly recruited to DNA damage sites and could not remain focused at micro-IR sites. The accumulation of XRCC1 ΔB2D progressively diffused and spread over time (Fig. 5C and 5F), and these defects were rescued by either GCN5 or TADA2B knockdown (Fig. 5F) but not by GCN5 inhibitor treatment (Suppl. Fig. 10C). We wondered whether diffused and unfocused accumulation of XRCC1 ΔB2D would be corrected at a later time point. ICC performed 15 minutes after micro-IR still showed diffuse and unfocused accumulation at micro-IR sites (Suppl. Fig. 10D). These data suggest that GCN5/TADA2B binding to XRCC1 is required for rapid and focused recruitment to DNA damage sites, which is critical for proper DNA damage repair. However, when XRCC1 has mutations in the BRCT II domain, GCN5/TADA2B binding becomes a constraining factor that prevents proper focused XRCC1 recruitment.

### GCN5/TADA2B provide dual regulatory control of XRCC1 recruitment

To investigate GCN5/TADA2B regulation of XRCC1 recruitment, we generated an *XRCC1* knockout U2OS cell line using CRISPR-Cas9 technology. This gene knockout did not alter GCN5 and TADA2B expression, but LIG3 protein was not detected, as XRCC1 is critical for its stability (Suppl. Fig. 11A). In *XRCC1* knockout U2OS cells, neither GCN5 nor TADA2B showed recruitment to micro-IR sites. However, reconstitution with XRCC1 in the knockout cells restored GCN5 and TADA2B recruitment to DNA damage sites, suggesting that GCN5/TADA2B protein binding to XRCC1 is not only essential for proper and focused recruitment of XRCC1, but also that GCN5 and TADA2B recruitment is XRCC1 dependent (Fig. 6A and 6B).

**Figure 6.**
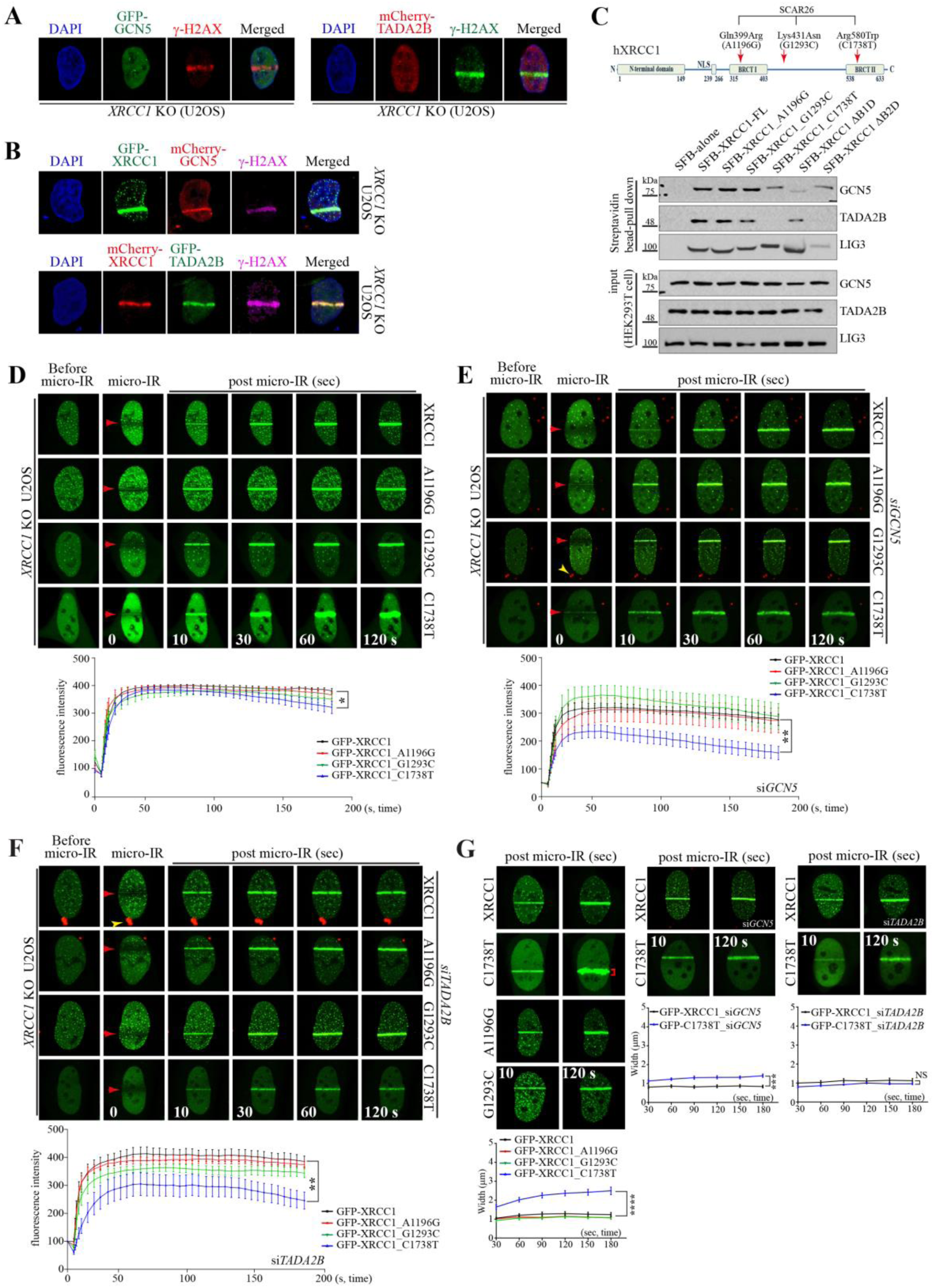
The XRCC1 mutant recruitment to DNA damage sites is regulated by GCN5 and TADA2B. **A.** GCN5 and TADA2B recruitment requires XRCC1. Immunocytochemistry analysis of *XRCC1* null U2OS cells expressing GFP-GCN5 and mCherry-TADA2B following micro-IR. γH2AX staining (Cy3 or FITC) marks damage sites. In the absence of *XRCC1*, neither GCN5 nor TADA2B accumulates at DNA damage sites, demonstrating XRCC1 dependent recruitment. **B.** XRCC1 reconstitution restores GCN5 and TADA2B recruitment. Rescue experiments in *XRCC1* null U2OS cells transfected with tagged XRCC1 (GFP or mCherry). γH2AX staining (Alexa Fluor 647) identifies micro-IR sites. XRCC1 expression fully restores the recruitment capacity of both GCN5 (mCherry) and TADA2B (GFP) to DNA damage sites. **C.** SCAR26 associated XRCC1 mutations show differential binding to GCN5 and TADA2B. Western blots of streptavidin pulldown assays with SFB tagged XRCC1 point mutants (c.1196A>G/p.Gln399Arg [referred to as A1196G], c.1293G>C/p.Lys431Asn [referred to as G1293C], c.1738C>T/p.Arg580Trp [referred to as C1738T]) demonstrate that Gln399Arg and Lys431Asn mutants retain binding to both GCN5 and TADA2B. However, the Arg580Trp mutant (located in BRCT II domain) maintains GCN5 and LIG3 binding but loses TADA2B interaction. XRCC1 deletion mutants (ΔB1D retains TADA2B binding and ΔB2D retains GCN5 binding) and LIG3 (which cannot bind ΔB2D) serve as controls. **D∼F.** Disease associated mutations, particularly C1738T, impair XRCC1 recruitment kinetics. Real-time imaging of GFP tagged WT and three point mutant XRCC1 (c.1196A>G [p.Gln399Arg], c.1293G>C [p.Lys431Asn], and c.1738C>T [p.Arg580Trp]) recruitments following micro-IR in control and gene depleted cells. Upper panels: Representative time course images showing GFP accumulation at damage sites (red arrowheads) over 180 seconds (s). Line graphs in lower panels: Quantitative recruitment kinetics. **D:** Comparison of WT versus three point mutants shows similar baseline recruitment, except C1738T. **E-F:** In *GCN5* (**E**) or *TADA2B* (**F**) depleted U2OS cells, the C1738T mutant (affecting BRCT II domain) exhibits severely impaired recruitment compared to WT and other point mutants. Gene depleted cells were identified by Cy3 labeled siRNA (yellow arrowheads). N=8-10 cells; **: p<0.01, *: p<0.05 **G.** The c.1738C>T mutation (C1738T) causes defective retention that is rescued by *GCN5/TADA2B* knockdown. Upper panels: Representative images showing GFP signal width over time. Line graphs in lower panel: Quantification of GFP signal spreading of WT and point mutant XRCC1. Analysis of protein spreading at damage sites reveals that the C1738T mutant shows progressive signal expansion (slope = 0.005), while other mutants remain focused like WT (slopes: WT = 0.001, A1196G = 0.000, G1293C = 0.001). *GCN5* or *TADA2B* knockdown restores focal accumulation of the C1738T mutant (slopes restored to control level [= 0.001] in all conditions), suggesting these factors contribute to the retention defect. N=7-10 cells; ****: p<0.001, ***: p<0.005, NS: not significant.

We also introduced three different point mutations (c.1196A>G, c.1293G>C, and c.1738C>T located in the BRCT I, CTL, and BRCT II domains, respectively) into the *XRCC1* construct, which are associated with SCAR26, and their sequences and expressions were verified (Fig. 6C, Suppl. Fig. 11B and 11C). When these constructs were introduced into *XRCC1* knockout U2OS cells, LIG3 proteins were restored. In addition, while XRCC1 ΔB1D introduction also restored LIG3 stability, the level of LIG3 was the same as in *XRCC1* knockout cells upon ΔB2D reconstitution, which lacks LIG3 binding sites (Suppl. Fig. 11D). SFB pulldown assay showed that A1196G and G1293C mutants could bind to GCN5, TADA2B, and LIG3. However, C1738T could bind to GCN5 and LIG3, but not TADA2B (Fig. 6C). This characteristic is similar to that of XRCC1 ΔB2D, which can bind to GCN5, but not TADA2B (Fig. 2B and 6C). Using real-time imaging, we examined the behavior of these three point mutant XRCC1 proteins. Overall recruitment of these three mutants to micro-IR sites was comparable to that of WT XRCC1; however, C1738T was not maintained at DNA damage sites over time (Fig. 6D). When either GCN5 or TADA2B was knocked down, recruitment of C1738T was significantly reduced (Fig. 6E and 6F), reminiscent of the recruitment behavior of XRCC1 ΔB2D at DNA damage sites (Fig. 5). Furthermore, C1738T could not remain focused at DNA damage sites, although the degree of diffusion was less than that of XRCC1 ΔB2D. The progressively diffused, unfocused retention of C1738T was corrected by GCN5 or TADA2B knockdown, following the same pattern as XRCC1 ΔB2D under either GCN5 or TADA2B knockdown conditions (Fig. 6G). These data suggest that GCN5 binding and its tethered TADA2B, which cannot bind to the mutant XRCC1 BRCT II domain, become a hindrance to focal retention of XRCC1 at DNA damage sites.

Collectively, our findings establish GCN5 and TADA2B as new critical regulators, independent of GCN5 HAT function, that operate through a dual mechanism: optimizing XRCC1 function under normal conditions while becoming counterproductive when XRCC1 structural integrity is compromised.

## DISCUSSION

DNA base damage, occurring thousands of times daily per cell, represents the most prevalent genomic insult. This damage is recognized and excised by specialized DNA glycosylases, generating apurinic/apyrimidinic (AP) sites that initiate the BER pathway. Subsequent processing by APE1 endonuclease creates single strand breaks within double stranded DNA structure, triggering SSBR mechanisms essential for genomic stability ^17–19,54,55^. The recruitment of APE1 and other critical BER/SSBR enzymes depends entirely on XRCC1, a scaffolding protein that orchestrates repair through extensive protein-protein interactions ^18,19^. Since the PARP1-XRCC1 protein interaction was established in the 1990s, XRCC1 recruitment to DNA damage sites has been understood to rely exclusively on PARP1 enzyme activation and subsequent PARylation, with PARP1 binding to the BRCT I domain proving essential for damage site localization ^18,55,56^, as we also demonstrated in the current study where XRCC1 lacking the BRCT I domain cannot be recruited to DNA damage sites. However, this PARP1 centric model has been insufficient to explain the precise spatial and temporal control of XRCC1 recruitment, particularly the mechanisms governing focal retention and maintenance at repair sites.

Here, we report the discovery of GCN5 and TADA2B — core and accessory components of the SAGA complex HAT module, respectively — as novel XRCC1 interacting proteins that expand our understanding of repair regulation. GCN5 and TADA2B interactions with XRCC1 operate through a constitutive binding mechanism independent of DNA damage status, distinct from conventional models of damage induced protein complex assembly. Our data demonstrate that these newly identified partners provide critical fine-tuning control over XRCC1 recruitment kinetics and focal retention at DNA damage sites, revealing additional regulatory layer that has remained hidden within the well-studied BER pathway.

These two proteins constitutively bind to XRCC1 regardless of DNA damage occurrence, and DNA damage does not further strengthen these protein interactions. Importantly, XRCC1 binding to GCN5 does not result in XRCC1 acetylation, despite GCN5’s established acetyltransferase function ^25^, regardless of DNA damage induction. Furthermore, pharmacological inhibition of GCN5 enzymatic activity did not alter XRCC1 recruitment kinetics to DNA damage sites, while GCN5 depletion significantly impaired recruitment. This finding suggests that the protein interaction itself, rather than enzymatic activity, represents the key functional feature, demonstrating a non-catalytic protein-enzyme interaction. Although GCN5 typically targets lysine (K) residues within KK, KXK, and EK amino acid motifs ^57^, comprehensive sequence analysis reveals no exclusive GCN5 acetylation sites within XRCC1. Database searches (PhosphoSitePlus, UniProt, CPLM: Compendium of Protein Lysine Modifications) identify potential target lysines including K260, K281, and K419; however, these sites are also targeted by other acetyltransferases such as PCAF, CBP/p300, and TIP60. Notably, SIRT1-mediated deacetylation of XRCC1 at K260, K298, and K431 prevents proteasomal degradation and stabilizes XRCC1 in chemoresistant lung cancer, though the responsible acetyltransferases remain to be identified ^58^. Additionally, PAR-binding screens identified XRCC1 but not GCN5 or TADA2B as PAR-binding proteins ^37^. However, GCN5 undergoes PARP1 mediated PARylation essential for recruitment to DSB sites, where it acetylates and crotonylates DNA-PKcs to facilitate repair ^59,60^. Moreover, GCN5 collaborates with E2F1 in NER ^61^. TADA2B’s involvement in these repair contexts has not been determined. In the current study, we demonstrate that GCN5 and TADA2B function as components of the SSBR machinery. We show that neither GCN5 nor TADA2B recruits to DNA damage sites in *XRCC1*-deficient cells, indicating that XRCC1 plays an essential role in their damage site localization and implying that recruitment of these three proteins to sites of DNA damage is mutually dependent. Furthermore, depletion of either GCN5 or TADA2B results in impaired DNA repair efficiency and increased cell death, demonstrating their functional importance in mammalian SSBR. While our data establish that XRCC1 is not a direct GCN5 substrate, we cannot exclude the possibility that GCN5 acetylates other BER/SSBR factors following XRCC1 mediated recruitment to damage sites. It has been reported that acetylation of BER/SSBR factors by p300 or CBP contributes to fine-tuning of repair efficiency ^62^, though GCN5’s involvement in this process remains to be explored. Additionally, GCN5 may facilitate repair through its well-established role in chromatin relaxation via histone modification at damage sites ^25^.

XRCC1 recruitment efficiency is significantly reduced when GCN5 and/or TADA2B are depleted, demonstrating that constitutive protein interactions among XRCC1, GCN5, and TADA2B are necessary for proper fine-tuning of XRCC1 recruitment. Since histones represent the primary targets of GCN5 enzymatic activity within the SAGA HAT module, the GCN5/TADA2B complex likely localizes near chromatin, poised to respond to stimuli ^26^. These interactions possibly position XRCC1 in proximity to DNA molecules, enabling rapid recruitment upon damage detection. This physiological configuration facilitates optimal and rapid XRCC1 recruitment to DNA damage sites. Our domain analysis reveals that both BRCT domains play distinct but essential roles in XRCC1 recruitment. The BRCT I domain, which serves as the primary GCN5 binding site, is absolutely critical for initial recruitment through PARP1 binding and PARylation responsiveness ^18,37^. Without the BRCT I domain, no subsequent BER or SSBR processes can occur. In contrast, the BRCT II domain, which binds TADA2B, is required for concentrated focal retention at damage sites. XRCC1 lacking the BRCT II domain (ΔB2D) exhibits compromised recruitment and progressive diffusion at damage sites, suggesting distinct functional roles for the two BRCT domains: BRCT I mediates recruitment itself, while BRCT II ensures concentrated retention, consistent with previous observations ^47^. Intriguingly, depletion of either GCN5 or TADA2B rescues the focal retention defects and diffused accumulation of XRCC1 ΔB2D, although recruitment kinetics show no improvement. This ΔB2D mutant retains the BRCT I domain that binds GCN5, which in turn maintains TADA2B tethering. This protein configuration may prevent the XRCC1 ΔB2D mutant from achieving optimal DNA proximity. By removing these protein constraints through GCN5/TADA2B depletion, focal retention of XRCC1 ΔB2D improves. These findings reveal that XRCC1 interactions with GCN5/TADA2B operate under dual regulatory control. Under normal conditions, GCN5 and TADA2B are essential for rapid recruitment and fine-tuning of XRCC1 accumulation at DNA damage sites. However, when XRCC1 is compromised — particularly through BRCT II domain mutations — these same protein interactions become inhibitory constraints that impede focused XRCC1 accumulation at damage sites. This represents a dynamic control mechanism where the regulatory network adapts its function based on XRCC1’s structural integrity.

To validate this regulatory model and explore its clinical relevance, we generated three SCAR26-associated XRCC1 point mutants. Overall recruitment kinetics of most mutants were comparable to WT XRCC1, except for the p.Arg580Trp (c.1738C>T) variant located in the BRCT II domain. This mutant, potentially relevant to SCAR26, retains LIG3 binding capacity but loses TADA2B interaction. Moreover, p.Arg580Trp exhibits defective focal accumulation and increased diffused accumulation at DNA damage sites, similar to but less severe than the XRCC1 ΔB2D phenotype. Importantly, this retention defect was rescued by GCN5 or TADA2B depletion, while recruitment kinetics deteriorated — mirroring the ΔB2D rescue pattern and reinforcing the critical role of GCN5/TADA2B in fine-tuning XRCC1’s rapid and focused recruitment. The similar behavior of XRCC1 ΔB2D and p.Arg580Trp — both retaining GCN5 binding but losing TADA2B interaction — demonstrates that disrupted XRCC1-TADA2B interactions create unfavorable protein configurations that impede focal accumulation and retention. Conversely, eliminating GCN5 or TADA2B restores favorable XRCC1 conformations for focal retention, albeit at the cost of rapid recruitment kinetics. These findings provide molecular insights into potential SCAR26 pathophysiology for future patient discovery.

Taken together as depicted in figure 7, we demonstrate that the GCN5/TADA2B complex serves as a novel fine-tuning regulator of XRCC1 recruitment to DNA damage sites, controlling both recruitment kinetics and focal retention. Unlike conventional damage induced repair protein interactions, the constitutive binding among these three proteins establishes a preparatory regulatory network that positions XRCC1 for rapid response upon damage detection. These findings reveal a previously unrecognized mechanism where chromatin associated factors pre-organize repair machinery to react readily upon DNA damage induction rather than responding reactively to damage signals. Importantly, our findings reveal the dual nature of this regulatory mechanism. While GCN5/TADA2B interactions optimize WT XRCC1 function, they become counterproductive when XRCC1 is structurally compromised. Specifically, BRCT II domain mutations convert beneficial regulatory interactions into inhibitory constraints that impede focal XRCC1 accumulation. This context dependent regulation explains how the same protein network can enhance normal repair while exacerbating dysfunction in disease states.

**Figure 7.**
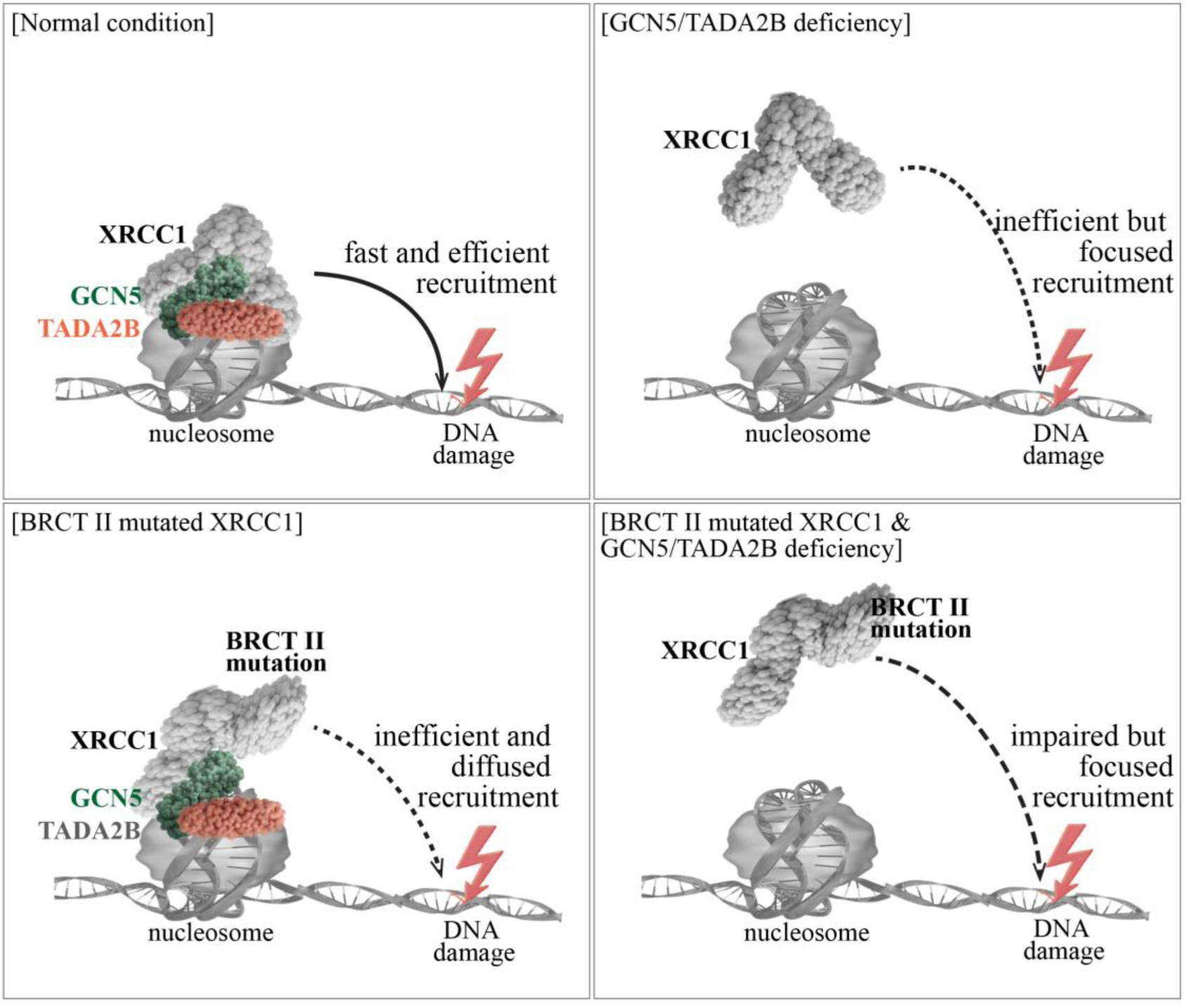
Schematic model of GCN5/TADA2B dual regulatory control of XRCC1 recruitment and retention at DNA damage sites. **(Normal condition)** Constitutive protein-protein interactions with GCN5/TADA2B position XRCC1 in proximity to DNA, leading to effective and focused recruitment to DNA damage sites. **(GCN5/TADA2B deficiency)** XRCC1 recruitment is impaired, but remains focused at DNA damage sites. **(BRCT II mutant XRCC1)** The mutant BRCT II domain cannot bind to TADA2B, leading to conformational disadvantage that results in diffused and ineffective recruitment to DNA damage sites. **(BRCT II mutant XRCC1 & GCN5/TADA2B deficiency)** The BRCT II mutant XRCC1 shows significantly impaired but focused recruitment to DNA damage sites.

## Supporting information

it contains all suppl. information

## ACKNOWLEDGEMENTS

We thank The Laboratory Animal Research Center (LARC) of Ajou University Medical Center for animal husbandry and the Three-Dimensional Immune System Imaging Core Facility at the Ajou University School of Medicine. We are grateful for technical supports from Jaemi Kim and Se-jeong Kim. YSL was supported by the NRF grant funded by the Korea government (MSIP) (RS-2017-NR021553 and RS-2022-NR069263).

## ROLE OF AUTHORS / CONTRIBUTIONS

YSL conceived the idea, and designed/supervised the project. KEK, JYK, DRL and YSL performed the experiments. MHK and BCY carried out the mass spectrometry analysis. JHJ provided critical reagents and helped interpretation of the data. All authors agree with the manuscript. YSL finalized the manuscript.

