## Supplementary material for "GCN5-TADA2B in the SAGA complex provide constitutive fine-tuning control of XRCC1 recruitment and focal retention at DNA damage sites": it contains all suppl. information

**Running title:** GCN5/TADA2B in XRCC1 machinery for DNA damage repair

7 Figures, 3 Supplementary Tables, 11 Supplementary Figures

**Supplementary table 1: PCR primer pairs for cloning**

| gene | Forward primer | Reverse primer | Reference |
|---|---|---|---|
| Mouse *Xrcc1* Full length | 5’-ATGCCGGAGATCAGCCTCCGC | 5’-GGCCTGGGGACCACCCCATA | Newly designed |
| All human genes below | | | |
| *XRCC1* Full length | 5’-ATGCCGGAGATCCGCCTCCG | 5’-GGCTTGCGGCACCACCCCATA | Newly designed |
| *XRCC1* NTD | 5’-CCGGAGATCCGCCTCCG | 5’-GTTGATCCGGCTGAAGA | Adapted from the article [1] |
| *XRCC1* NTL | 5’-TTCCGTGTGAAGGA | 5’-GGTGCCTTCTCCTCGGG | |
| *XRCC1* B1D | 5’-CCAGCTCCAACTCGTACC | 5’-ATCCTCCTCACTGCTGGA | |
| *XRCC1* CTL | 5’-ATGGCAGGGCCAGGTTC | 5’-CTCAGGGACTGGCAGATC | |
| *XRCC1* B2D | 5’-CAGGAGCCTCCTGATCTG | 5’-GGCTTGCGGCACCACCCC | |
| **Stepwise cloning strategy to generate a *XRCC1* B1D or B2D domain deletion mutants.** | | | |
| *XRCC1* ΔB1D | Step1: *XRCC1* Full length forward primer | Step1:5’-GGAACCTGGCCCTGCCTCCTCTGGGCCAGC | Newly designed |
| | Step2:5’-GCTGGCCCAGAGGAGGCAGGGCCAGGTTCC | Step2: *XRCC1* Full length reverse | |
| | Step3: *XRCC1* full length forward for PCR products Steps 1/2 | Step3: *XRCC1* full length reverse for PCR products Steps 1/2 | |
| *XRCC1* ΔB2D | *XRCC1* Full length forward | *XRCC1* CTL reverse | |
| **Point mutation *XRCC1* cloning primers (the mutants were selected on the website for ClinVar: https://www.ncbi.nlm.nih.gov/clinvar/)** | | | |
| *XRCC1*_ A1196G in the 1st BRCT domain | 5’-CGGCTGCCCTCCCGGAGGTACCTCATG | 5’-CATGAGGTACCTCCGGGAGGGCAGCCG | Newly designed |
| *XRCC1*_ G1293C in the CTL domain | 5’-AAGCTTCCTCAGAACCAACCCCAGACCAAA | 5’-TTTGGTCTGGGGTTGGTTCTGAGGAAGCTT | |

| | | | |
|---|---|---|---|
| *XRCC1_* C1738T in the 2nd BRCT domain | 5'-GAGGACTATATGAGTGACTGGGTTCAGTTTGTGATC | 5'-GATCACAAACTGAACCCAGTCACTCATATAGTCCTC | |
| *TADA2B* Full length | 5'-GCGGAGCTGGGGAAGAA | 5'-AGACGCGTCCCTGGAGATCG | Newly designed |
| *TADA2B* Zn/SANT | 5'-GCGGAGCTGGGGAAGAA | 5'-CCCCAGGTTCCCGTGGAT | Adapted from the article [2] |
| *TADA2B* CTL | 5'-GGGAAGGCCTGCATCC | 5'-AAGGTTCTCAATGGCGG | |
| *TADA2B* SWIRM | 5'-CTTCCAGGCTTCGAGCT | 5'-AGACGCGTCCCTGGAGAT | |
| *GCN5* Full length | 5'-GCGGAACCTTCCCAGGC | 5'-CTTGTCAATGAGGCCTCCCTC | Newly designed |
| *GCN5* NTD | 5'-GCGGAACCTTCCCAGGC | 5'-GCGCTCCTCCAGGCGGG | Adapted from the article [3] |
| *GCN5* ACT | 5'-CCTATGCCAGGCGAGAAG | 5'-CTTGTCAATGAGGCCTCC | |
| *GCN5* BROMO | 5'-CTGATGGAGTGTGAGCTGA | 5'-CTTGTCAATGAGGCCTCC | |

**Supplementary table 2-1: siRNA sequences to knock down the target gene expression**

| gene | Sense Sequence | Antisense sequence |
|---|---|---|
| *XRCC1* siRNA | 5’ GGAAACUCAUCCGAUACGUUU 3’ | 5’ ACGUAUCGGAUGAGUUUCCUU 3’ |
| *TADA2B* siRNA | 5’ CAUUACGUGAGCAUGUACAUU 3’ | 5’ UGUACAUGCUCACGUAAUGUU 3’ |
| *GCN5* siRNA | 5' GAAGCUGAUUGAGCGCAAA 3' | 5' UUUGCGCUCAAUCAGCUUC 3' |

The primers were designed using the following websites.
http://sidirect2.rnai.jp/
https://eurofinsgenomics.eu/
https://rnaidesigner.thermofisher.com/rnaiexpress/

**Supplementary table 2-2: CRISPR primer sequences to knock out the target gene expression**

| gene | Sense Sequence | Antisense sequence |
|---|---|---|
| *XRCC1* | 5’CACCGTCTCCGGCATGTCAACGTCG | 5’AAACCGACGTTGACATGCCGGAGAC |

The primers were designed using the following websites.
https://chopchop.cbu.uib.no/
https://www.milliporesigmabioinfo.com/bioinfo_tools/faces/informatics.xhtml

**Supplementary table 3: List of antibodies for Western blot and Immunocytochemistry**

| Antibody | Company | Raised in | Cat # | Titer |
|---|---|---|---|---|
| XRCC1 | Abcam | Rabbit | Ab9147 | 1:2,000 |
| LIG3 | BD biosciences | Mouse | #611876 | 1:2,000 |
| PNKP | Abcam | Rabbit | Ab181107 | 1:2,000 |
| POLB | Abcam | Rabbit | Ab26343 | 1:500 |
| TADA2B | Abnova | Mouse | H00093624 | 1:1,000 |
| GCN5 | Cell signaling Technology | Rabbit | #3305 | 1:2,000 |
| GST | Thermo Fisher Scientific | Mouse | MA4-004 | 1:2,000 |
| GFP | Santa Cruz Biotechonology | Mouse | SC-9996 | 1:2,000 |
| FLAG | Santa Cruz Biotechonology | Mouse | SC-166355 | 1:5,000 |
| mCherry | Invitrogen | Rabbit | PA5-34974 | 1:2000 |
| ATM | Abcam | Rabbit | Ab199726 | 1:2,000 |
| ATM-pS (serine)1981 | Cell signaling Technology | Rabbit | #5883 | 1:2,000 |
| P53 | Abcam | Mouse | Ab28 | 1:5,000 |
| P53-pS15 | Cell signaling Technology | Rabbit | #9284 | 1:1,000 |
| CHK1 | Abcam | Rabbit | Ab32531 | 1:1,000 |
| CHK1-pS345 | Cell signaling Technology | Rabbit | #2348 | 1:1,000 |
| CHK2 | Millipore | Mouse | #05-649 | 1:1,000 |
| CHK2-pT(threonine)68 | Novus Biologicals | Rabbit | NBP3-13076 | 1:2,000 |
| KAP1 | Novus Biologicals | Rabbit | NBP2-76411 | 1:5,000 |
| KAP1-pS824 | Novus Biologicals | Rabbit | NBP2-76398 | 1:2,000 |
| H2AX | Cell signaling Technology | Rabbit | #7631 | 1:2,000 |
| γ-H2AX | Cell signaling Technology | Rabbit | #2577 | WB1:2,000<br>ICC 1:500 |
| Histone 3 (H3) | Abcam | Rabbit | Ab1791 | 1:50,000 |
| K-acetylation | Abcam | Rabbit | Ab190479 | 1:1,000 |
| Actin | Origene Technologies | Mouse | TA811000 | 1:2,000 |
| tubulin | Personal gift<br>[Dr. Sang Gyu Park] | Mouse | N/A | 1:40,000 |

### SUPPLEMENTARY FIGURE LEGENDS

**Supplementary Figure 1. Confirmation of GST fused protein expression and effective knockdown of gene expression by siRNA.**

**A.** Validation of GST fusion human protein expression of XRCC1, GCN5, and TADA2B. Left panel: Ponceau S staining demonstrates overall protein expression levels in bacterial lysates expressing GST fused human XRCC1, GCN5, and TADA2B (pDEST15 vector). Middle panel: Anti GST Western blot confirms successful pulldown of GST fusion proteins from bacterial lysates. Right panel: Western blot analysis using protein specific antibodies validates the identity of purified GST fusion proteins. Arrowheads indicate bands of interest.

**B.** Validation of SFB tagged protein expression in mammalian (HEK293T) cells. Western blot analysis using anti FLAG antibodies confirms expression of SFB tagged human XRCC1, GCN5, and TADA2B constructs following streptavidin pulldown from transfected cell lysates. The SFB system incorporates FLAG epitopes enabling detection of fusion proteins. Arrowheads indicate successfully expressed SFB tagged target proteins.

**C.** Confirmation of efficient gene knockdown by siRNA in HeLa and U2OS cells. Western blot analysis demonstrates effective protein depletion using gene specific siRNAs. Two independent siRNA sequences were tested for *TADA2B* knockdown and both siRNA worked equally well; siRNA #1 was selected for subsequent experiments. Notably, *GCN5* knockdown also reduces TADA2B protein levels (arrowheads), indicating interdependent protein stability.

**D.** TADA2B stability depends on GCN5 expression. Western blot analysis shows treatment with the proteasome inhibitor MG132 partially restores TADA2B protein levels in *GCN5* knockdown cells, demonstrating that TADA2B stability is GCN5 dependent. This finding parallels the well characterized XRCC1-LIG3 stability relationship (Figure 1B).

**Supplementary Figure 2. Expression confirmation of truncated proteins.**

**A~C.** Validation of truncated protein constructs. Western blot analysis confirms expression of deletion mutants for domain mapping studies: (A) five XRCC1 truncation mutants, (B) three GCN5 truncation mutants, and (C) three TADA2B truncation mutants, all fused with GST or SFB tags. Left panels: Schematic diagrams showing the specific domains retained in each truncation construct with corresponding amino acid positions and predicted molecular weights. Right panels: Western blot detection using GST or FLAG antibodies demonstrates expression at expected molecular weights (red arrowheads), confirming successful generation of all truncation mutants for binding domain analysis.

**D.** Expression validation of XRCC1 BRCT domain deletion mutants. Western blot confirmation of two additional XRCC1 constructs lacking either BRCT I domain (ΔB1D) or BRCT II domain (ΔB2D). These deletion mutants were specifically designed to assess the individual contributions of each BRCT domain to protein-protein interactions. Red arrowheads indicate successfully expressed proteins at predicted molecular weights.

**Supplementary Figure 3. Prediction and validation of protein interaction among XRCC1, GCN5, and TADA2B.**

**A.** Cell-free validation of BRCT domain binding specificity. GST pulldown assays using immobilized truncated XRCC1 proteins confirm the domain mapping results from Figure 2A. Western blot analysis demonstrates that both BRCT I and BRCT II domains are necessary for optimal GCN5 and TADA2B binding in a reconstituted system, validating the cellular interaction data.

**B.** Mapping of GCN5 interaction domains. SFB pulldown assays identify the minimal GCN5 regions required for XRCC1 and TADA2B binding. Upper panel: Schematic representation of GCN5 truncation mutants showing domain boundaries and amino acid positions. Lower panel: Western blot analysis reveals that the ACT part is primarily responsible for XRCC1 interaction, while additional regions contribute to TADA2B binding.

**C.** Mapping of TADA2B interaction domains. SFB pulldown assays define the TADA2B regions mediating XRCC1 and GCN5 interactions. Upper panel: Schematic diagram of TADA2B truncation constructs with domain annotations. Lower panel: Western blot analysis identifies the CTL portion as the interaction interface for both XRCC1 and GCN5.

**D.** Computational prediction supports experimental binding data. AlphaFold structural modeling of the XRCC1-GCN5-TADA2B complex provides theoretical support for the observed interactions.

Upper panel: Domain architecture of XRCC1 comparison showing experimental truncation boundaries relative to predicted structural domains from The Encyclopedia of Domains (TED). Green boxes indicate predicted aligned error regions of XRCC1, GCN5, and TADA2B. Lower panel: Two different versions of AlphaFold prediction were used to foresee the interactions of three proteins [4,5]. AlphaFold 3 prediction[5] of the trimeric complex (pTM = 0.43) shows GCN5 positioned near BRCT I and TADA2B spanning both BRCT domains, consistent with biochemical data. This confidence score compares favorably to the established XRCC1-LIG3 interaction (pTM = 0.33, data not shown), supporting the biological relevance of these novel protein interactions.

**Supplementary Figure 4. Protein interactions remain unaltered before and after exposure to DNA damaging reagents.**

**A-D.** Constitutive binding is maintained following DNA damage induction. SFB pulldown assays in HeLa cells demonstrate that XRCC1, GCN5, and TADA2B interactions are unaffected by DNA damage treatment. Cells were exposed to different DNA damaging agents for 2 hours: (A) methyl methanesulfonate (MMS, 0.4 mM) - alkylating agent causing base modifications and replication fork stalling; (B) camptothecin (CPT, 2 μM) - topoisomerase I inhibitor inducing replication dependent DNA strand breaks; (C) phleomycin (Phleo, 20 μg/ml) - radiomimetic agent causing direct DNA strand breaks; and (D) hydrogen peroxide ($H_2O_2$, 500 μM) - oxidizing agent generating base lesions and DNA strand breaks. Western blot analysis reveals equivalent binding intensities before (untreated) and after drug treatment, confirming that these protein interactions are constitutively maintained rather than damage induced, distinguishing them from typical DNA damage response protein complexes.

**Supplementary Figure 5. Confirmation of cloned gene expression.**

**A.** Denaturing protocol for XRCC1 acetylation analysis. Schematic diagram illustrates the methodology for detecting XRCC1 lysine (K) acetylation using denaturing conditions followed by SFB pulldown. This approach removes all protein-protein interactions prior to pulldown, ensuring that detected acetylation signals originate solely from XRCC1 rather than co-precipitating acetylated proteins.

**B.** Validation of GFP tagged protein expression. Western blot analysis confirms successful expression of GFP fused constructs (pDEST53 vector) in transfected HEK293T cells. Protein expression was verified using antibodies against GFP, XRCC1, GCN5, and TADA2B. Both exogenous (GFP-tagged) and endogenous proteins are detected, with red arrowheads indicating endogenous protein levels for comparison. Expression levels demonstrate successful overexpression.

**C.** Validation of mCherry tagged protein expression of XRCC1, GCN5, and TADA2B. Western blot confirmation of mCherry fused protein constructs demonstrates successful expression.

**D.** Expression validation of GFP or mCherry tagged BRCT deletion mutants. Western blot analysis confirms expression of XRCC1 constructs lacking either BRCT I (ΔB1D) or BRCT II (ΔB2D) domains, tagged with either GFP or mCherry fluorescent proteins. Arrowheads indicate successfully expressed deletion mutants migrating at predicted molecular weights. Right panel: Schematic representation of ΔB1D and ΔB2D domain architectures.

**Supplementary Figure 6. Gain of function analysis by XRCC1, GCN5, or TADA2B overexpression for cell viability.**

**A.** Overexpression does not confer protection against DNA damage. Colony formation assays comparing control (CTRL) and gene overexpressing U2OS cells (*XRCC1*, *GCN5*, or *TADA2B*) following continuous exposure to DNA damaging agents for 9-10 days: phleomycin (Phleo), methyl methanesulfonate (MMS), hydrogen peroxide ($H_2O_2$), or camptothecin (CPT) at indicated concentrations. Left panels: Representative Crystal Violet staining demonstrates similar colony formation patterns across all conditions. Line graphs in right panels: Quantitative survival analysis reveals no significant protective effect from protein overexpression, contrasting with the clear sensitization observed upon gene depletion (Figure 3A). N=6.

**B.** Long term viability analysis confirms lack of protective effect. Sulforhodamine B (SRB) assays measuring cell viability in control versus gene overexpressing U2OS cells treated with DNA damaging agents at indicated time points. Line graphs in left panels: Growth curves show comparable proliferation rates between control and overexpressing cells before and after exposure to DNA damaging reagents. Bar graphs in right panels: Relative viability at day 6 demonstrates no significant differences compared to untreated controls, confirming that increased protein levels do not enhance DNA damage tolerance. N=6; NS: not significant.

**C.** Cell cycle responses remain normal in overexpressing cells. FACS analysis of cell cycle distribution in control and gene overexpressing U2OS cells following 24 hour DNA damage treatment. Histograms in left panels: Representative cell cycle profiles show appropriate checkpoint activation across all conditions. Bar graphs in right panels: Quantification of G1, S, and G2/M phase distributions reveals normal cell cycle responses, indicating that overexpression does not disrupt checkpoint signaling or enhance recovery from DNA damage induced arrest. N=6.

**D.** Gene depletion reduces basal proliferation rates. SRB proliferation assays comparing control and gene depleted U2OS cells (*XRCC1*, *GCN5*, or *TADA2B* siRNA) under normal growth conditions without cellular stress. Time course analysis demonstrates that protein depletion significantly impairs cell proliferation even in the absence of exogenous DNA damage, suggesting these proteins are necessary for normal cellular homeostasis. N=6; ***: $p<0.005$, *: $p<0.05$.

**Supplementary Figure 7. Annexin V and propidium iodide (PI) FACS analysis to detect cell death.**

**A~E.** Representative dot plots of Annexin V/PI flow cytometry. Annexin V positive/PI negative cells were considered early stage apoptotic, and Annexin V/PI double positive cells were considered late stage apoptotic. (A) No treatment, (B) phleomycin (Phleo), (C) methyl methanesulfonate (MMS), (D) hydrogen peroxide ($H_2O_2$), and camptothecin (CPT) treated U2OS cells are shown. The conditions for drug exposure are indicated in the figure. The numbers indicate a representative percentage of either Annexin V positive (right lower quadrant) or Annexin V/PI double positive (right upper quadrant) cells.

**Supplementary Figure 8. Gain of function analysis by *XRCC1, GCN5,* or *TADA2B* overexpression for DNA damage responses.**

**A~D.** Overexpression does not enhance DNA damage response signaling. Western blot analysis of DNA damage response pathways in control (CTRL) and gene overexpressing U2OS cells (*XRCC1*, *GCN5*, or *TADA2B* using pDEST53 vector) following treatment with: (A) phleomycin (20 μg/ml, 1 hour), (B) methyl methanesulfonate (2 mM, 1 hour), (C) hydrogen peroxide (500 μM, 30 minutes), or (D) camptothecin (2 μM, 1 hour). Cells were analyzed immediately after treatment (0h) and at 3 hours post-treatment (3h). All overexpressing cell lines demonstrate normal activation of DNA damage response markers (ATM phosphorylation, γ-H2AX formation, checkpoint signaling) comparable to control cells, with appropriate kinetics of response resolution. NT: no treatment.

**Supplementary Figure 9. γH2AX foci resolution is delayed in *XRCC1, GCN5,* and *TADA2B* knockdown U2OS cells compared to control cells after DNA damage (expanded data of Figure 4E).**

**A-D.** Quantitative analysis of DNA repair kinetics using γH2AX foci resolution. Immunocytochemistry analysis of control and gene depleted U2OS cells (*XRCC1*, *GCN5*, or *TADA2B* siRNA) following brief DNA damage pulses to assess repair efficiency over 24 hours. DNA damage was induced for 30 minutes with (A) phleomycin (20 μg/ml), (B) methyl methanesulfonate (1 mM), or (D) camptothecin (1 μM), or for 10 minutes with (C) hydrogen peroxide (500 μM). Left panels: Representative images showing γH2AX foci (Cy3 immunofluorescence) at key time points: no treatment (NT), immediately post-damage (30 min or 10 min), 6 hours, and 24 hours after damage removal. All knockdown cell lines (*XRCC1*, *GCN5*, *TADA2B*) show normal initial foci formation but significantly impaired foci resolution compared to controls. Bar graphs in right panels: Quantitative analysis demonstrates persistent γH2AX signal in gene depleted cells at three time points, indicating incomplete DNA repair. N>300 nuclei per condition; ****: p<0.001, ***: p<0.005, **: p<0.01, *: p<0.05.

**Supplementary Figure 10. Defective recruitment of XRCC1 to DNA damage sites in absence of *GCN5* or *TADA2B*.**

**A.** GCN5 enzymatic activity is not required for XRCC1 recruitment. Real-time analysis of XRCC1 recruitment to micro-irradiated sites in U2OS cells treated with or without GCN5 inhibitor (GCN5i, Butyrolactone 3). Upper panel: Representative images showing comparable XRCC1 accumulation at DNA damage sites regardless of GCN5 inhibition. Red arrowheads indicate the sites of micro-irradiation. Lower panel: Quantitative kinetic analysis demonstrates no significant difference in recruitment efficiency, confirming that GCN5's acetyltransferase activity is dispensable for XRCC1 localization. N=9-10 cells; NS: not significant.

**B.** Defective recruitment of BRCT mutant XRCC1 is not affected by GCN5 enzyme activity. Live-cell imaging of GFP tagged XRCC1 constructs (wildtype, ΔB1D, ΔB2D) in GCN5 inhibitor (GCN5i, Butyrolactone 3) treated cells following micro-irradiation. Upper panel: Representative time course images (0-120 seconds) showing GFP accumulation at damage sites (red arrowheads indicate micro-irradiation sites). Lower panel: Recruitment kinetics over 180 seconds reveal that XRCC1 ΔB1D fails to recruit, while ΔB2D shows severely reduced recruitment efficiency. These results parallel those in Figure 5B, confirming that BRCT domain requirements are independent of GCN5 enzymatic function. N=9-10 cells; ***: $p<0.005$, **: $p<0.01$, *: $p<0.05$.

**C.** BRCT II deletion impairs focal accumulation and retention independent of GCN5 enzyme activity. Analysis of GFP signal width at DNA damage sites in GCN5 inhibitor treated cells demonstrates that XRCC1 ΔB2D exhibits both reduced recruitment and defective focal retention, appearing as broadened signal distribution. This phenotype mirrors the findings in Figure 5F, further supporting that GCN5's scaffolding function, rather than its enzymatic activity, is critical for proper XRCC1 positioning, focal accumulation, and retention. Upper panels: Representative images showing GFP signal width (red bracket) over time. Line graphs in lower panels: Quantification of signal spreading (both wildtype and ΔB2D slope = 0.001). . N=9-10 cells; ***: $p<0.005$.

**D.** Validation of XRCC1 ΔB2D recruitment defects by immunocytochemistry. Fixed cell analysis of GFP tagged XRCC1 constructs (wildtype and ΔB2D) and γH2AX at 1 and 15 minutes post micro-irradiation confirms the live-cell imaging results. GFP signals (from tagged constructs) and γH2AX immunostaining (Alexa Fluor 647) demonstrate the persistent recruitment and retention defects of BRCT deletion mutants in a fixed-cell format.

**Supplementary Figure 11. Successful generation of *XRCC1* knockout cell and XRCC1 point mutants.**

**A.** Validation of *XRCC1* knockout U2OS cell line. Western blot analysis confirms complete loss of XRCC1 protein expression following CRISPR-Cas9 gene editing. As expected, LIG3 protein levels are also reduced due to the established XRCC1-dependent stability relationship. Importantly, GCN5 and TADA2B expression remains normal, confirming that *XRCC1* deficiency does not affect these proteins and validating this knockout cell line for rescue experiments.

**B.** Sequence verification of SCAR26 associated XRCC1 point mutations. DNA sequencing confirmation of successfully generated point mutants corresponding to disease associated variants: c.1196A>G (A1196G/p.Gln399Arg, referred to as A1196G), c.1293G>C (G1293C/p.Lys431Asn, referred to as G1293C), and c.1738C>T (C1738T/p.Arg580Trp, referred to as C1738T). Red circles indicate the targeted nucleotide changes, confirming precise genome editing.

**C.** Expression validation of XRCC1 point mutant constructs. Western blot analysis using anti-GFP (upper panel) and anti-XRCC1 (lower panel) antibodies confirms successful expression of all mutant constructs. Red arrows indicate endogenous XRCC1 expression.

**D.** Functional reconstitution of *XRCC1* knockout cells. Western blot analysis of *XRCC1*-null U2OS cells transfected with wildtype or three point mutant XRCC1 constructs demonstrates successful XRCC1 protein expression. LIG3 levels are restored in all XRCC1 expressing cell lines including XRCC1_ΔB1D, yet not in ΔB2D deletion mutant, which lacks the BRCT II domain required for LIG3 binding and stabilization. GCN5 and TADA2B expression remains consistent across all conditions. Of note, Anti-XRCC1 antibody cannot detect XRCC1 ΔB2D.

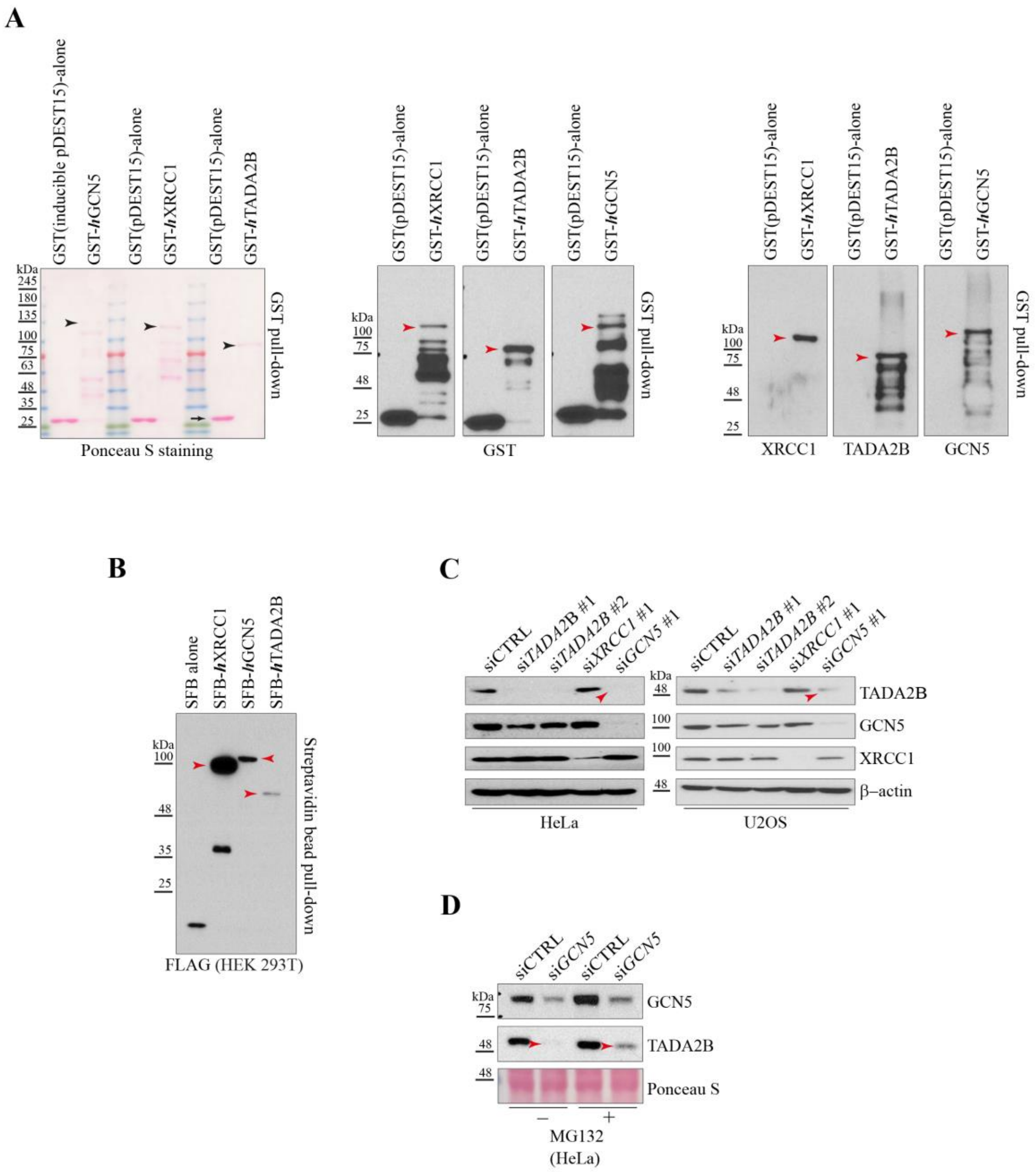
A
GST(inducible pDEST15)-alone
GST-hGCN5
GST(pDEST15)-alone
GST-hXRCC1
GST-hTADA2B
GST pull-down
Ponceau S staining
GST
XRCC1
TADA2B
GCN5
kDa
B
SFB alone
SFB-hXRCC1
SFB-hGCN5
SFB-hTADA2B
Streptavidin bead pull-down
FLAG (HEK 293T)
C
siCTRL
siTADA2B #1
siTADA2B #2
siXRCC1 #1
siGCN5 #1
β–actin
HeLa
U2OS
D
siGCN5
Ponceau S
MG132
(HeLa)

Suppl. Fig. 2 (Tada2b)

A

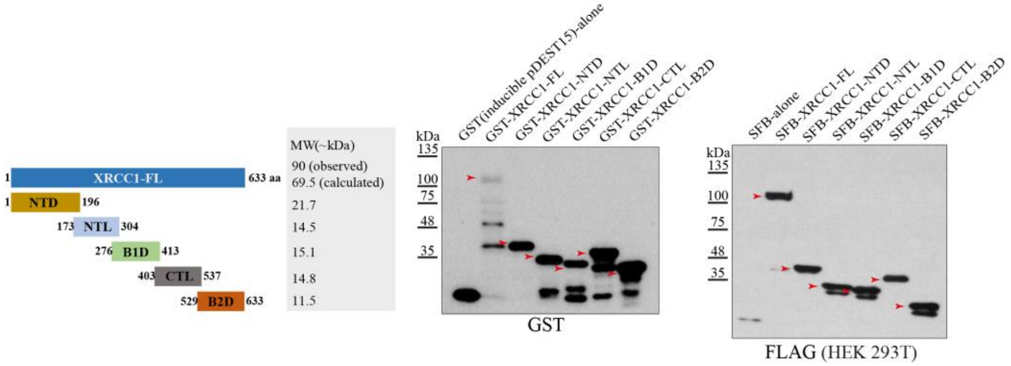

B

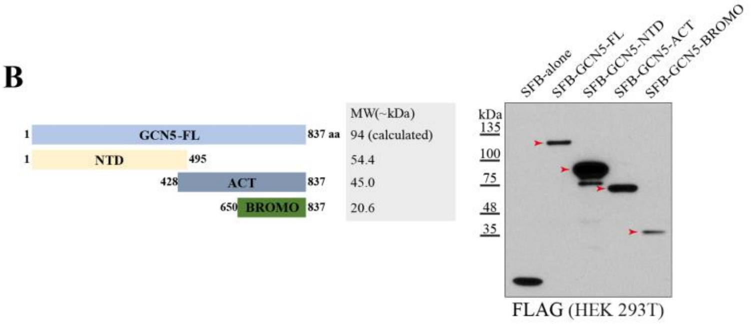

C

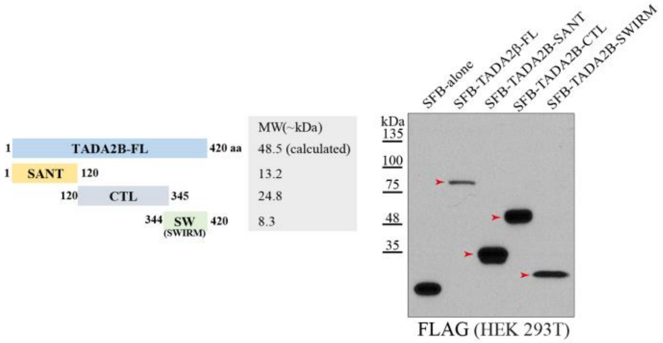

D

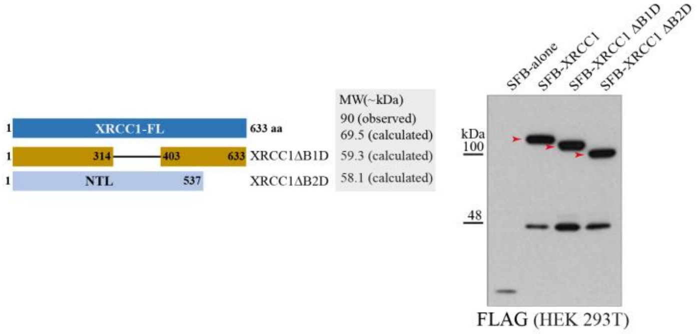

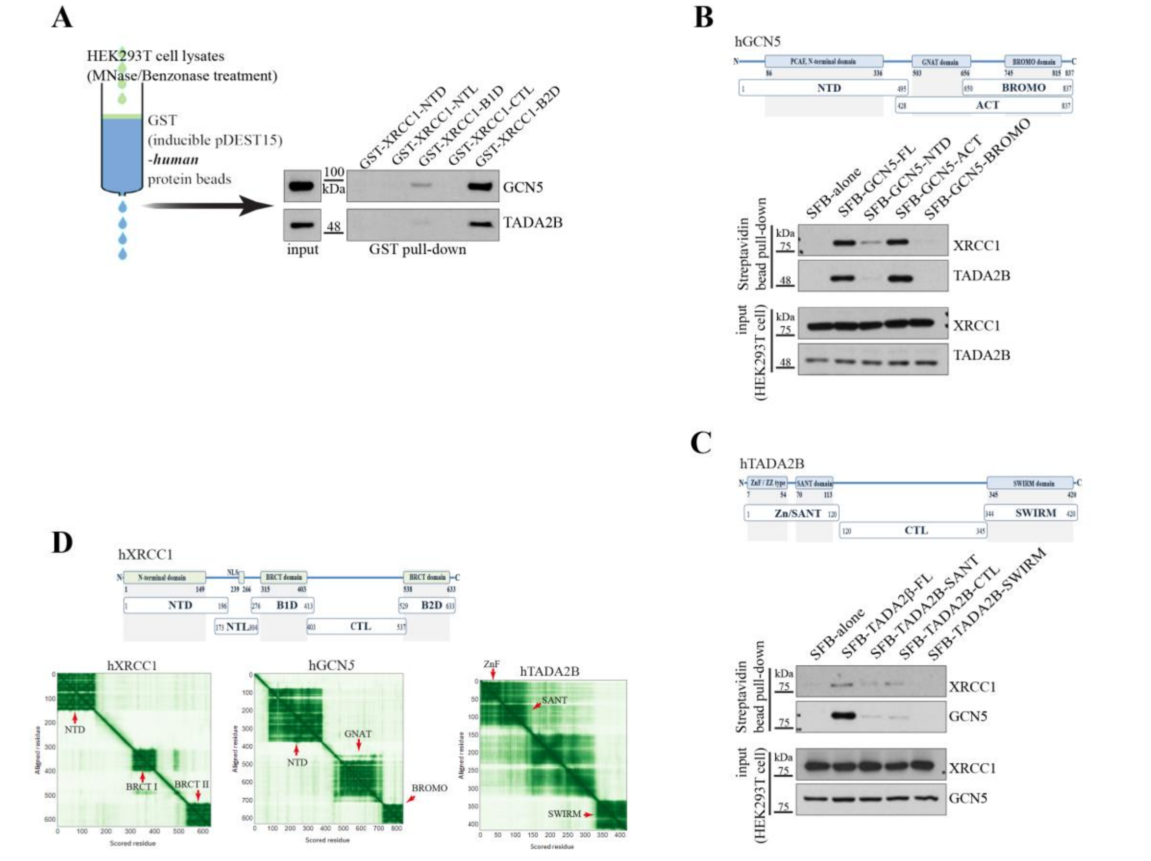

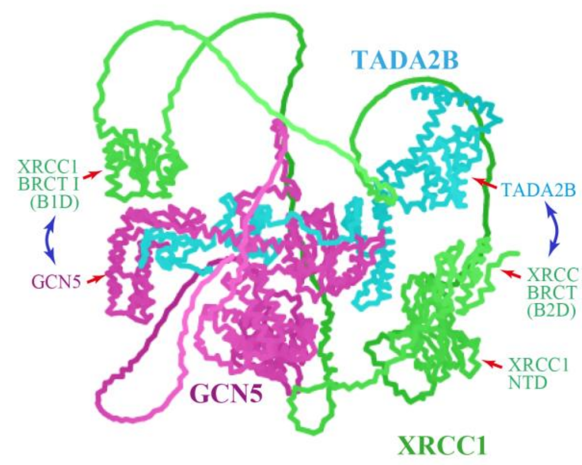

*ColabFold: making protein folding accessible to all ; Nature Methods, 19, 679-682 (2022)*

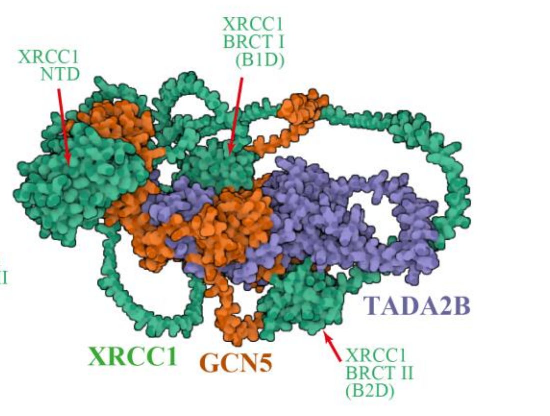

*Accurate structure prediction of biomolecular interactions with AlphaFold3;Nature, 630, 493-500 (2024)*

Suppl. Fig. 4 (Tada2b)

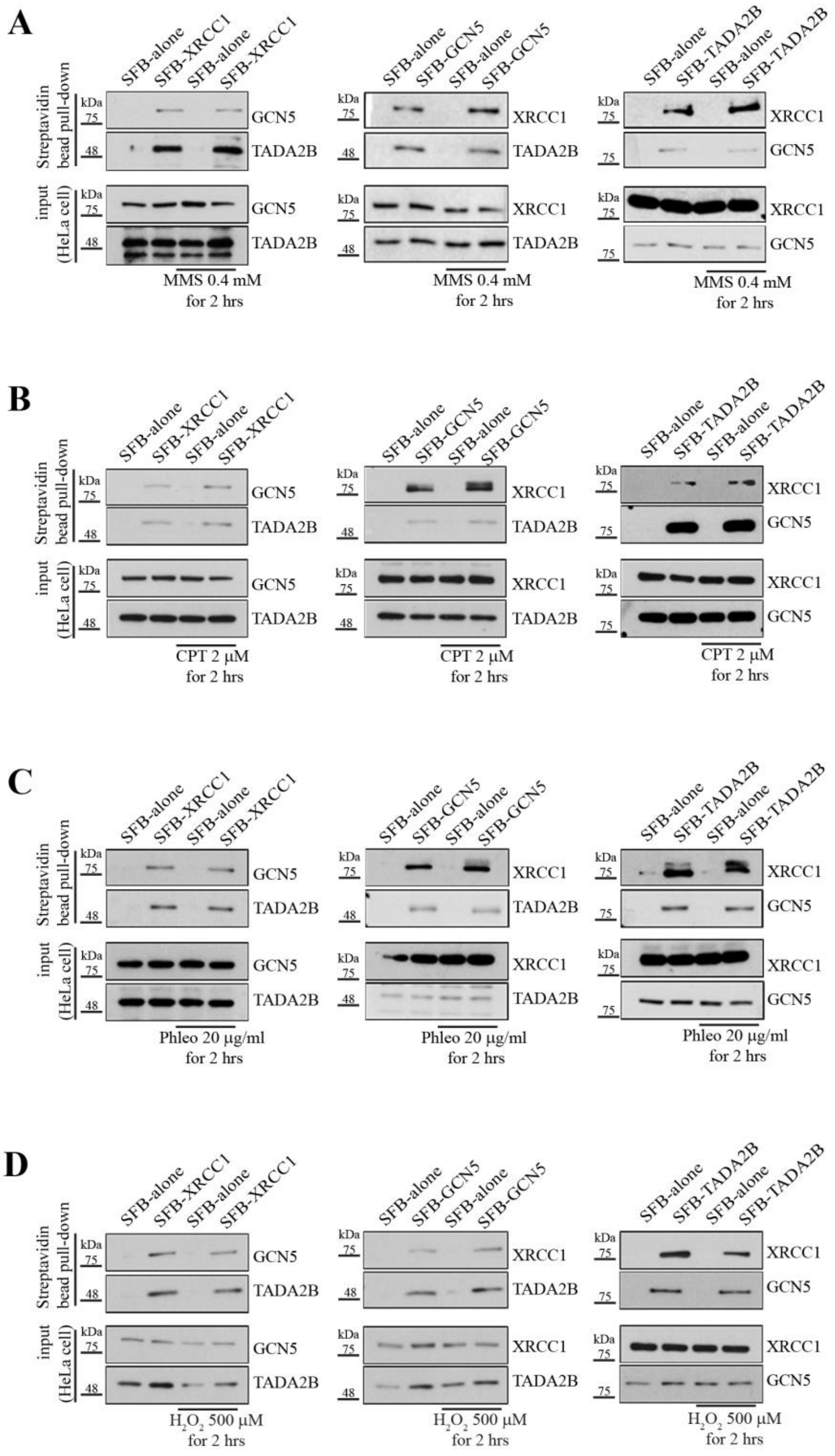

Suppl. Fig. 5 (Tada2b)

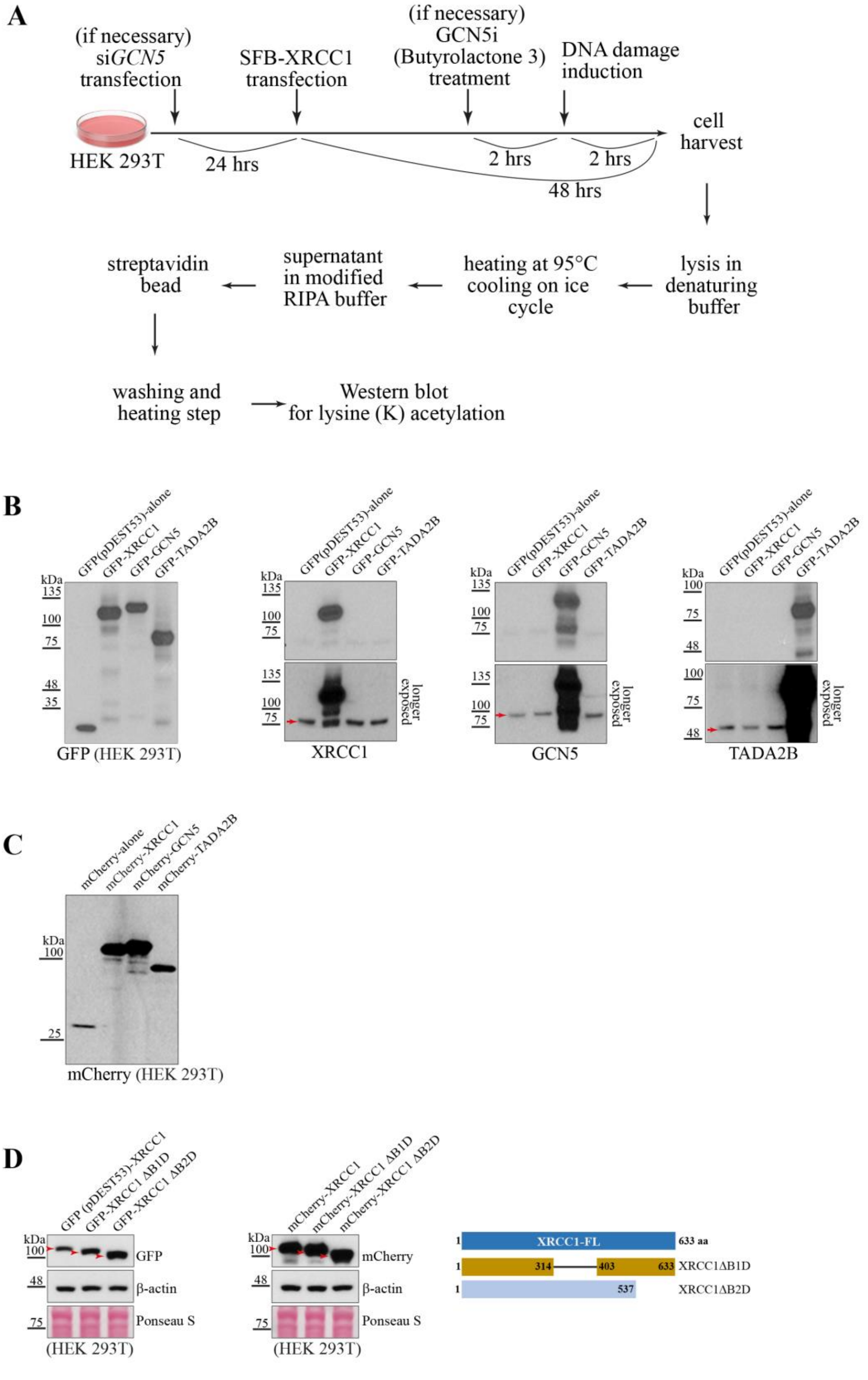

Suppl. Fig. 6 (Tada2b)

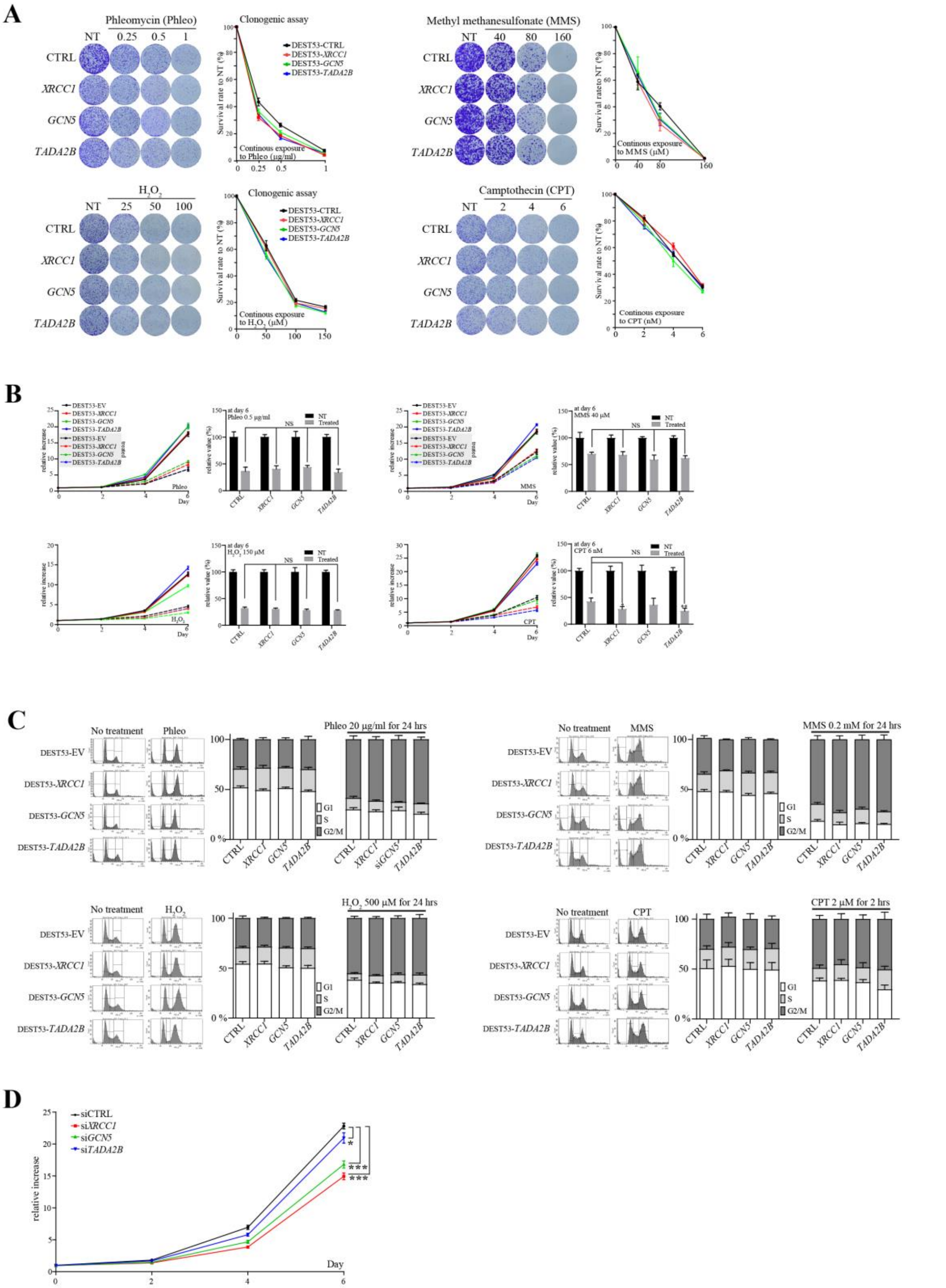

Suppl. Fig. 7 (Tada2b)

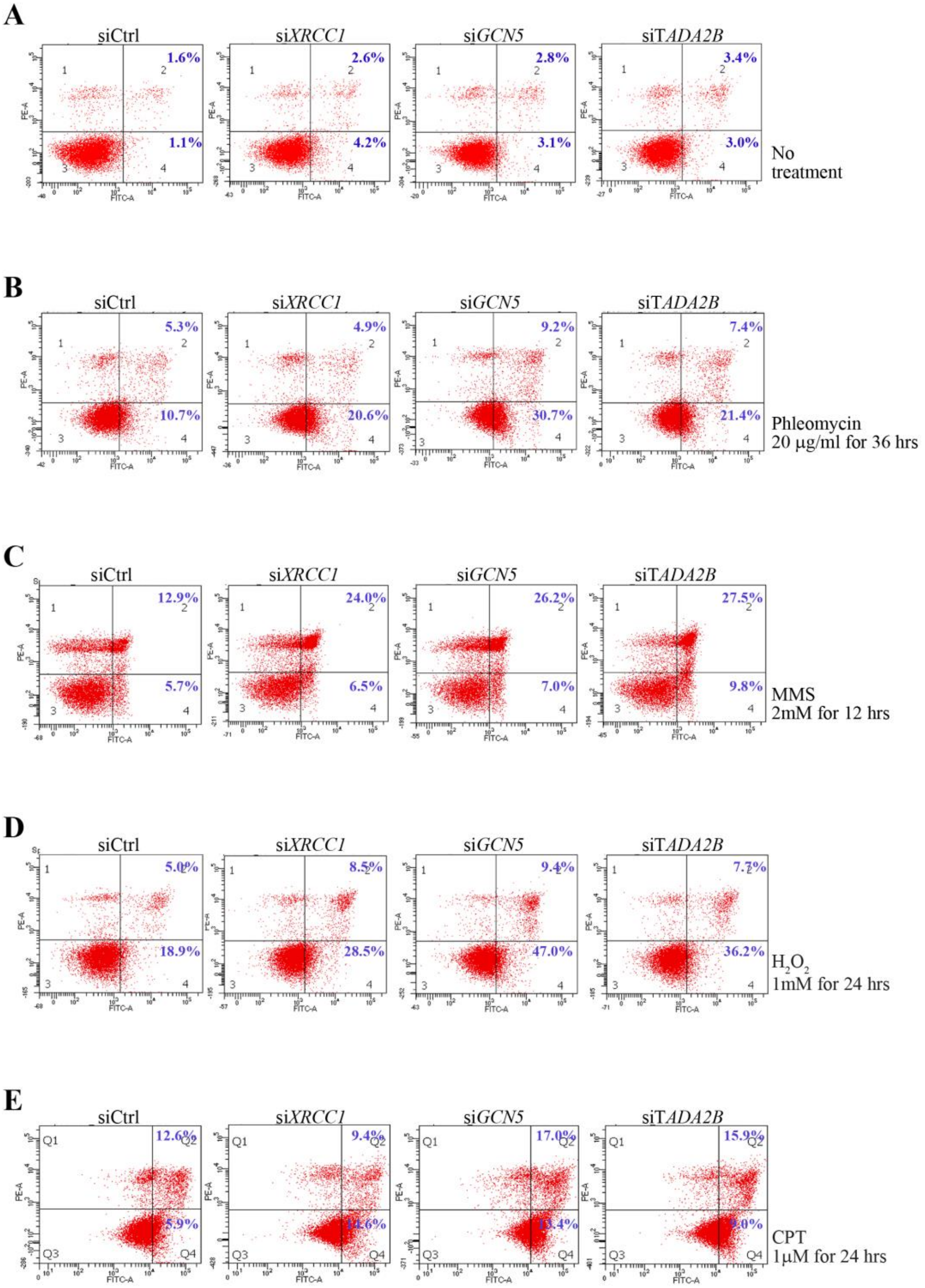

Suppl. Fig. 8 (Tada2b)

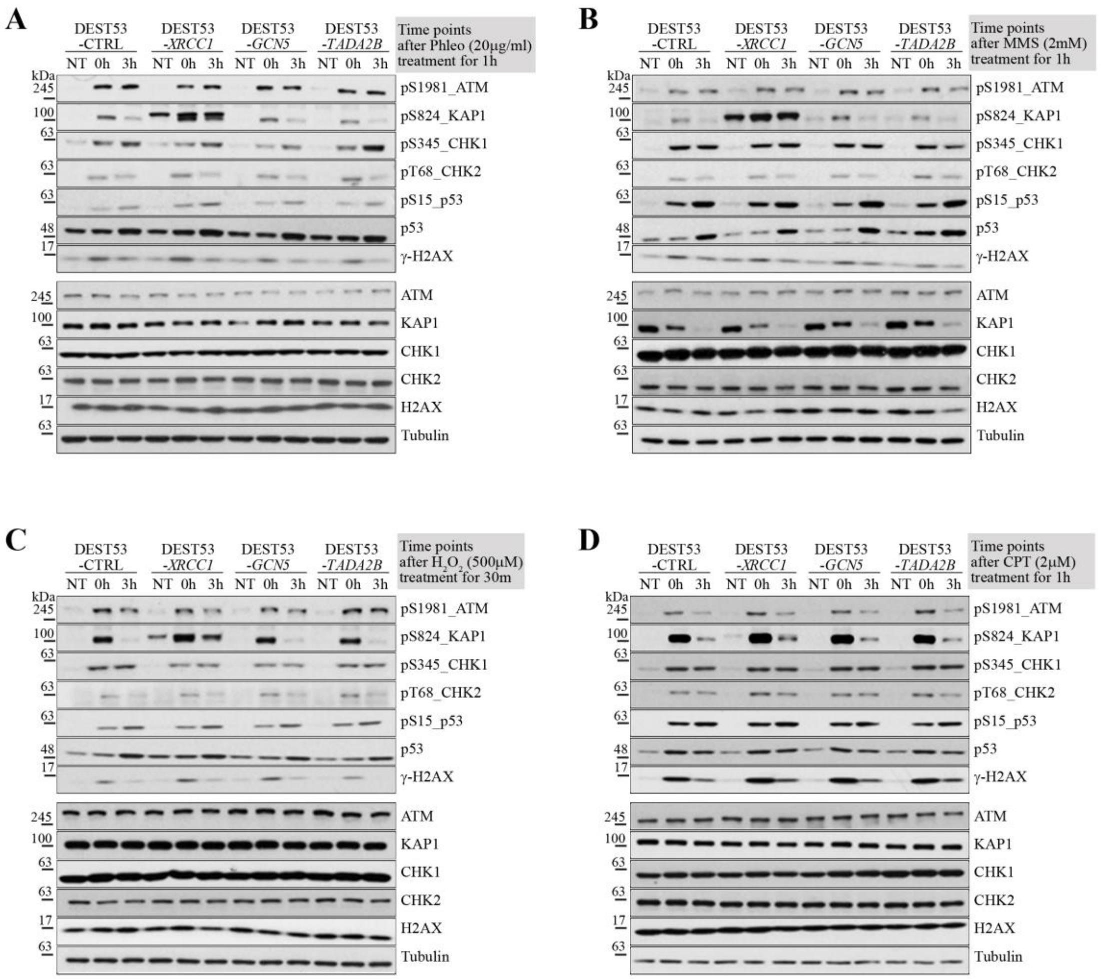

Suppl. Fig. 9 (Tada2b)

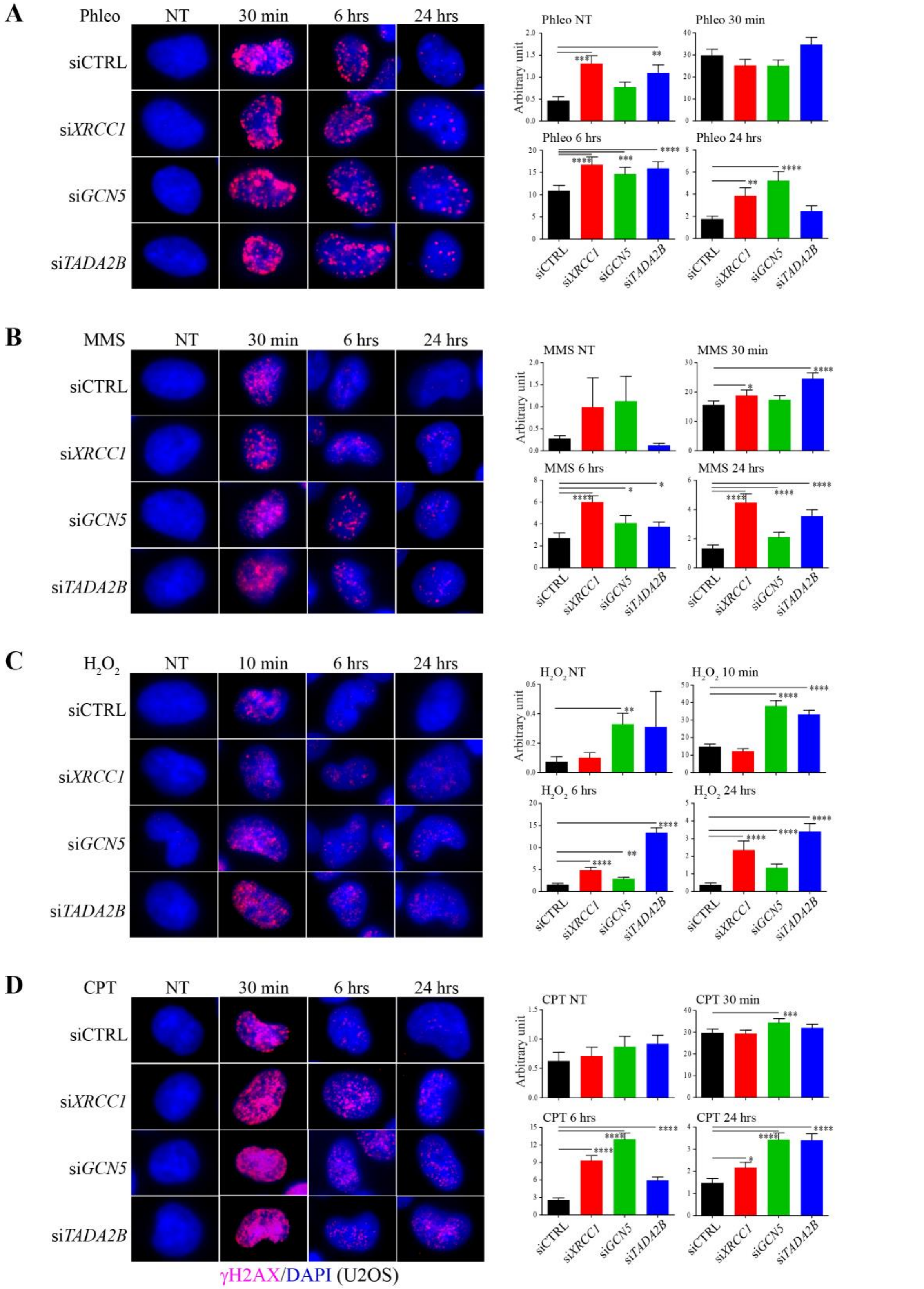

Suppl. Fig. 10 (Tada2b)

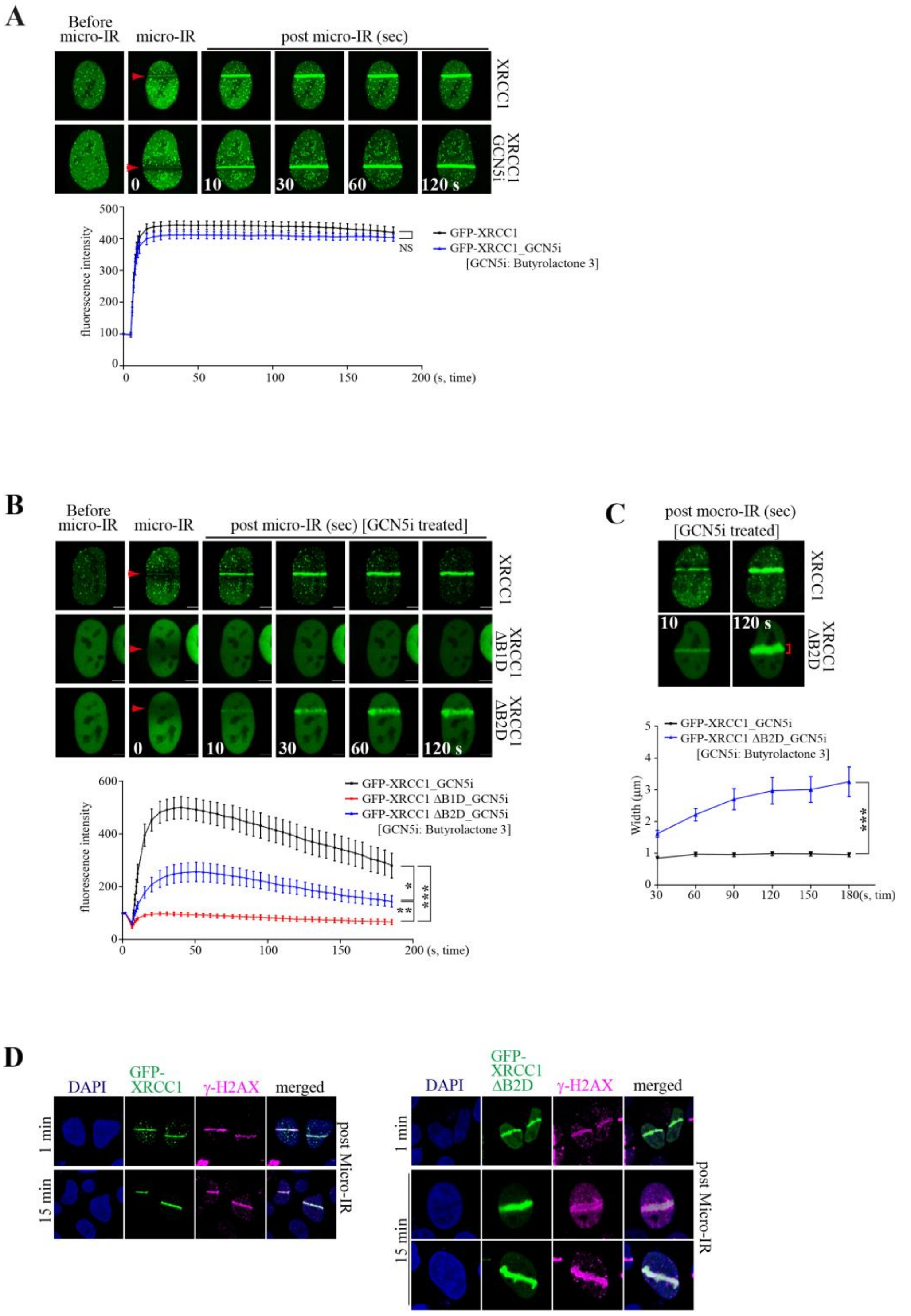

Suppl. Fig. 11 (Tada2b)

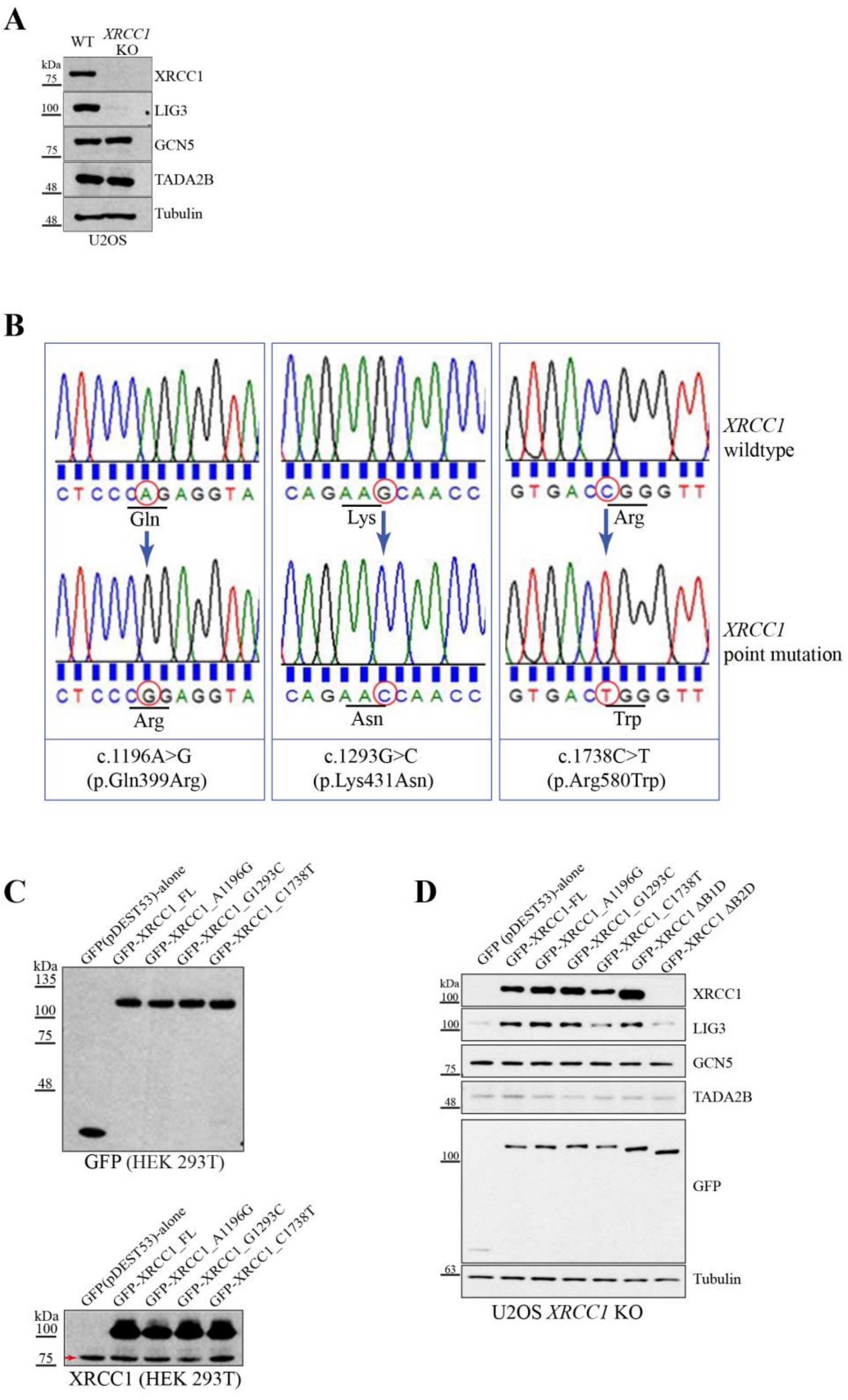
